# Cerebral vasculature shapes the spatial patterning of brain metastases

**DOI:** 10.64898/2026.07.22.739598

**Authors:** Asa Farahani, Zhen-Qi Liu, Vincent Bazinet, Charles P. Couturier, Evan Porter, Dante P. I. Capaldi, Olivier Morin, Alain Dagher, Bratislav Misic

## Abstract

Primary cancer cells that originate in diverse tissues in the body can spread to the brain through various physiological pathways and form metastatic tumors. The spatial patterning of brain metastases is highly stereotyped across individuals, but the physiological factors that confer regional vulnerability to tumor colonization are poorly understood. Here we map the brain metastases associated with primary breast cancer, lung cancer, and malignant melanoma to the multiscale organization of the brain. We analyze structural magnetic resonance imaging (MRI) data from more than 2, 300 cancer patients with approximately 10, 000 brain metastases to estimate metastatic frequency maps. We then examine whether transcriptional (microarray profiling), functional (functional MRI), and vascular (arterial spin labeling) features can predict the spatial pattern of each brain metastasis type. Our analyses highlight vascular anatomy as an integral determinant of metastatic patterning, particularly in breast and lung cancers. We find arterial border-zones as sites of elevated vulnerability to metastatic invasion. These areas, characterized by slow flow and small-caliber vessels, create a hemodynamic environment that favors the arrest and extravasation of circulating tumor cells. Together, these results provide quantitative support for the long-standing hypothesis that vascular architecture and its biomechanical properties constrain metastatic seeding in the brain. Identifying vascular topology as a key determinant of metastatic patterning suggests that systemic vascular and metabolic factors may contribute to metastatic risk.

## INTRODUCTION

Brain metastases occur with varying frequency across different primary cancer types^69,107^. Breast cancer, lung cancer, and melanoma are among the malignancies with the highest propensity for brain metastasis^80,106^. Importantly, metastatic tumors are not uniformly distributed across the brain. This spatial selectivity is conceptualized by the “fertile soil” hypothesis, which posits that specific brain microenvironments are preferentially permissive to metastatic colonization^22,87^. This permissiveness depends on regional differences in brain metabolic activity, gene expression profiles, and vascular architecture, as well as on the physical and molecular properties of circulating tumor cells originating from the primary tumor^6,23,35,73^. Understanding the factors that render certain brain regions more susceptible to tumor colonization provides insights for developing more effective and targeted prevention and therapeutic strategies.

Advances in neuroimaging and large-scale open-access data initiatives have enabled a suite of new integrative investigations in cancer studies. A recent study by Barrios and colleagues provided brain metastasis frequency maps for breast cancer, lung cancer, and melanoma, quantifying how often metastasis is localized in a brain region across a large cohort of patients^8^. Concurrently, extensive datasets such as the Human Connectome Project (HCP)^14,42,43,50,119^ and the Allen Human Brain Atlas (AHBA)^52^ have enabled the construction of normative brain atlases that feature functional, vascular, and transcriptomic properties of brain tissue. These atlases can further be used to estimate how similar biological profiles are across brain regions and to construct inter-regional similarity matrices where each edge encodes the pairwise similarity between two regions with respect to a given biological feature^9,49^. Integrating tumor frequency maps with inter-regional similarity matrices allows us to test whether regions with similar biological profiles also exhibit similar susceptibility to metastatic involvement, and provides a framework to identify the factors that shape region-specific metastatic vulnerability in the brain.

Here we construct whole-brain inter-regional similarity matrices spanning architectural, hemodynamic, vascular, and transcriptomic domains, and ask which biological similarity matrix explains the spatial distribution of brain metastases across cancer types. We find that vascular similarity between regions explains the patterning of metastasis frequency in breast and lung cancers. Furthermore, we show that the normative arterial transit time (ATT) map is the biological feature that is positively associated with brain metastasis frequency maps in the primary breast and lung cancers. This finding provides quantitative support for prior qualitative observations that implicate the so-called “watershed” regions as sites of elevated risk for metastatic cell colonization. Given prior evidence that aging and adverse metabolic health negatively affect vascular dynamics, particularly in arterial border-zones^32,33^, we propose that systemic metabolic factors should be incorporated into studies of brain metastasis and into the assessment of patient-level brain metastatic risk.

## RESULTS

Brain metastasis frequency maps were obtained from the study by Barrios and colleagues^8^ through the Open Science Framework (OSF) at https://osf.io/fkqmr. These maps show the spatial distribution of metastatic tumor localization, and reflect how frequently different brain regions are affected across patients. The frequency map for primary breast cancer was estimated from 573 participants with a total number of 3, 078 lesions (Fig. 1a), the frequency map for primary lung cancer was estimated from 1, 323 participants with a total number of 5, 324 lesions (Fig. 1b), and the frequency map for primary melanoma was estimated from 409 participants with a total number of 1, 996 lesions (Fig. 1c). Cohort characteristics, including age, biological sex, and tumor histology, are summarized in Table. S1. Full methodological details are provided in the original publication^8^ and are summarized in *Methods*.

**Figure 1.**
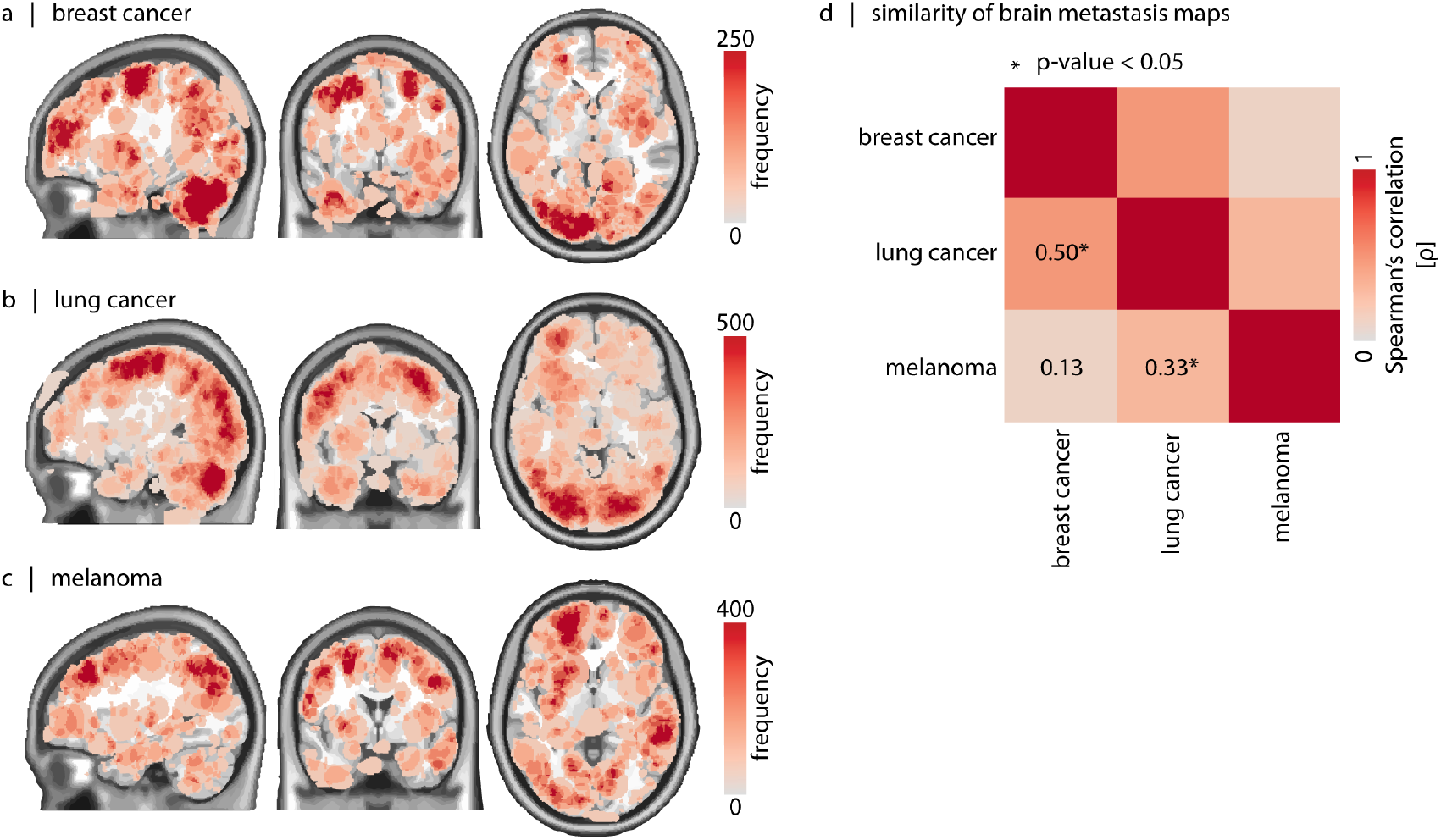
Brain metastasis frequency maps. Brain metastasis frequency maps are shown on sagittal, coronal, and axial views of the MNI152 template for patients with (a) primary breast cancer (*N* = 573 patients, *L* = 3, 078 lesions), (b) primary lung cancer (*N* = 1, 323 patients, *L* = 5, 324 lesions), and (c) melanoma (*N* = 409 patients, *L* = 1, 996 lesions). Brain maps are sourced from Barrios and colleagues^8^. Data are shown on a T1-weighted group average template (MNI152). (d) Similarity between brain metastasis frequency maps, quantified using Spearman’s rank correlation coefficient (*ρ*). Correlations are computed after parcellating the metastasis maps at the level of cortex (Schaefer-400^105^) and cerebellum (probabilistic atlas of the cerebellar lobules^24^). Asterisks (*) indicate statistical significance based on spatial autocorrelation-preserving null models (FDR-corrected *p*_moran_ < 0.05). We further show that the parcellated cancer-specific metastasis frequency maps deviate from a volume-proportional spatial distribution. For each cancer type, we generate 1, 000 null maps using a multinomial model in which regional sampling probabilities are proportional to cerebellar and cortical parcel volume, while total metastasis burden is preserved. We quantify deviation from the null model using Σ _*i*_ (*d*_*i*_ − *E*_*i*_)^2^*/E*_*i*_, where *d*_*i*_ and *E*_*i*_ are the observed and expected metastasis burdens in region *i*. All three cancer-specific maps deviate from their null models (*p* = 9.99 × 10^−4^), indicating regional predilections for metastasis beyond parcel-volume effects.

The pseudo-continuous arterial spin labeling (ASL) and resting-state functional MRI data were obtained from the HCP Lifespan studies (2.0 Release)^14,50^. ASL data were used to estimate maps of arterial transit time (ATT) and cerebral blood flow (CBF). ATT quantifies the time required for magnetically labeled blood to reach different brain parts from the labeling plane (cervical region) and CBF quantifies the amount of regional blood perfusion (mL/100g/min). The functional MRI timeseries were used to estimate functional connectomes. The microarray gene expression data were derived from the Allen Human Brain Atlas (AHBA)^52^. See *Methods* for more details.

### Unique and divergent patterns of metastasis frequency maps

Metastatic tumors are not distributed uniformly across the brain. Each primary cancer type exhibits a distinct spatial pattern of brain metastasis frequency. Notably, the distribution patterns for breast and lung metastases are more similar to one another (*ρ* = 0.50, false discovery rate (FDR)-corrected *p*_moran_ = 5.49 × 10^−3^) than to melanoma (breast–melanoma: *ρ* = 0.33, FDR-corrected *p*_moran_ = 2.75 × 10^−2^; lung–melanoma: *ρ* = 0.13, FDR-corrected *p*_moran_ = 0.65) (*N*_moran_ = 1, 000) (Fig. 1). Breast and lung cancers preferentially metastasize to the cerebellum and posterior-parietal cortex, whereas melanoma metastases are less concentrated in the cerebellum. These observations are consistent with the notion that cancer cells invade a permissive “fertile soil” that supports their growth, and do not invade all brain regions in the same manner. Tissue permissiveness is shaped by both cancer-specific biological properties, such as the originating cell type and associated molecular adhesion mechanisms, and intrinsic features of the host tissue microenvironment.

To assess the robustness of the spatial pattern of metastasis frequency, we compared the metastasis frequency maps for breast and lung cancers (shown in Fig. 1a,b^8^) with independent frequency maps published by Bao and colleagues^6^, available at https://www.synapse.org/#!Synapse:syn57077557/files. Despite differences in cohort size (177 patients with breast cancer brain metastases and 548 with lung cancer brain metastases^6^), the corresponding maps have consistent spatial organization. The Spearman’s rank correlation between frequency maps for metastatic breast cancer is *ρ* = 0.48 (*p*_moran_ = 1.99 × 10^−3^, *N*_moran_ = 1, 000), and the correlation between frequency maps for metastatic lung cancer is *ρ* = 0.56 (*p*_moran_ = 9.99 × 10^−4^, *N*_moran_ = 1, 000). This concordance indicates that the observed patterns are not specific to a single cohort but instead reflect reproducible patterns of metastatic spread in the brain (Fig. S1). In the following subsection, we seek to identify the biological basis that shapes the organization of the metastasis frequency maps and ask which forms of brain interregional biological similarity explain regional susceptibility to metastatic colonization.

### Mapping metastatic burden to multi-scale biological networks

We next evaluate whether the spatial distribution of metastatic burden is organized according to homophilic principles in the brain’s intrinsic biological architecture. Metastatic burden is defined as the frequency of metastatic lesions within each brain region. Rather than testing whether metastasis simply co-localizes with the spatial pattern of a biological feature, we test whether regions embedded in similar biological neighbourhoods exhibit similar susceptibility to metastatic involvement. In this context, neighbourhoods are not restricted to physical adjacency. The same brain region can have different neighbours depending on whether inter-regional proximity is defined by spatial distance, functional connectivity, vascular organization, or transcriptomic similarity. To test this idea, we construct multiple interregional similarity networks spanning spatial, functional, vascular, and molecular domains (Fig. 2a), and examine which network can explain the spatial organization of metastatic burden. Specifically, we include a distance network, capturing the Euclidean proximity between regions; a functional connectivity network, capturing similarity in spontaneous hemodynamic fluctuations between regions; vascular similarity networks, capturing inter-regional co-variation in cerebral blood flow (CBF) and arterial transit time (ATT) across subjects, reflecting shared vascular architecture (see *Methods* for further details); and a transcriptomic similarity network, capturing similarity in regional gene-expression profiles. Collectively, these analyses assess whether metastatic vulnerability is embedded within any of the mentioned intrinsic similarity architectures of the brain.

**Figure 2.**
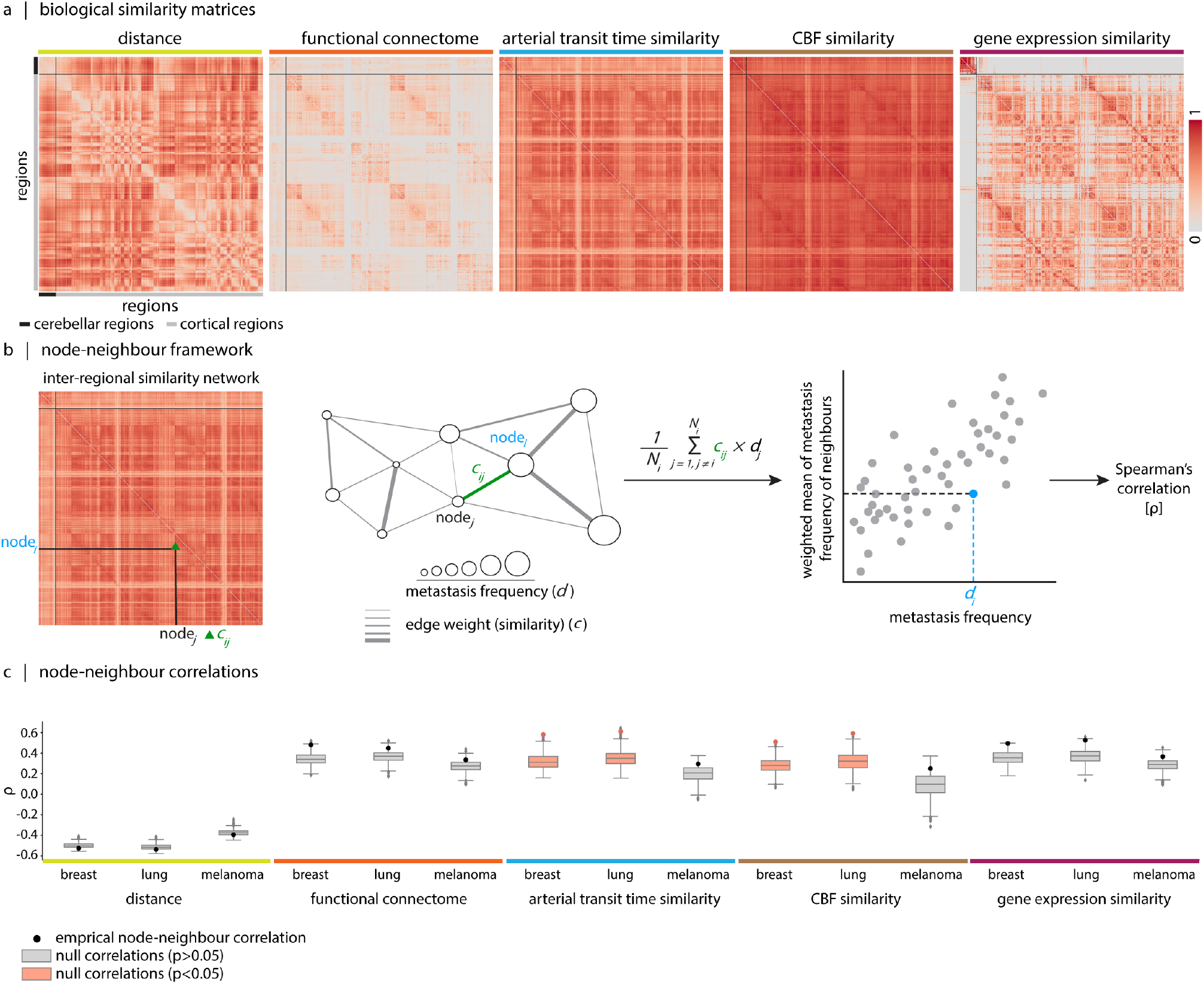
Brain metastasis frequency maps for breast and lung cancer are shaped by vascular network similarity. We examine how brain inter-regional biological similarity networks shape the spatial organization of brain metastasis frequency maps. (a) For each biological brain feature, a region × region similarity network is constructed, where each edge represents how close or similar two regions are with respect to that feature. Heatmaps depict Euclidean distance, hemodynamic, arterial transit time, cerebral blood flow, and transcriptomic similarity matrices. Only positive edges are shown. These matrices include both cerebellar and cortical regions, given their relevance to metastatic burden (Fig. 1a–c). (b) Node-neighbour similarity framework. The metastatic exposure of region_i_ is defined as the weighted average metastatic burden of its neighbours, where weights correspond to the strength of connections in the similarity network of interest. Formally, exposure is computed as the mean of *c*_*ij*_ *× d*_*j*_ across all regions *j* with positive connectivity to region_i_, where *c*_*ij*_ denotes the inter-regional similarity (edge weight) and *d*_*j*_ denotes the metastatic burden of region_j_. We then correlate each region’s metastatic burden (*d*_*i*_) with its metastatic exposure to assess whether regions connected to highly affected neighbours also exhibit greater metastatic burden. (c) Empirical correlations (*ρ*) between regional metastatic burden and neighbour-weighted burden, shown alongside null distribution of correlations. Boxplots in orange indicate statistical significance based on spatial autocorrelation-preserving null models (FDR-corrected *p*_moran_ < 0.05). Notably, the distance matrix is included as a control, as spatially autocorrelated null models are not expected to yield significant associations under this framework. In Fig. S3, we limit the node-neighbour framework to cortical regions and further extend the analysis to include metabolic, receptor, and laminar similarity matrices.

To assess the contribution of each inter-regional similarity network (Fig. 2a) to the spatial patterning of metastatic frequency, we adopt a “node-neighbour” similarity assessment framework^31,49,1^ . We define the metastatic exposure of region_i_ to the rest of the brain as the weighted average metastatic burden of its neighbouring regions, with weights given by the strength of connections in the similarity network of interest. Formally, exposure is computed as the mean of *c*_ij_ *× d*_j_ across all regions *j* with positive connectivity to region_i_, where *c*_ij_ denotes inter-regional similarity (edge weight) and *d*_j_ denotes the metastatic burden of region_j_. We then correlate each region’s metastatic burden (*d*_i_) with its metastatic exposure to test whether regions strongly connected to highly affected neighbours also exhibit greater metastatic burden. A positive correlation indicates that the spatial pattern of metastasis aligns with the underlying inter-regional similarity architecture (see Fig. 2b). To ensure that the observed correlation is specifically driven by network topology rather than spatial autocorrelation of the metastatic frequency map, we generate spatially constrained null frequency maps (*N*_moran_ = 1, 000) and assess the statistical significance of the empirical correlation relative to the corresponding null correlation distribution.

For primary breast and lung cancers, metastasis frequency maps are primarily linked to the architecture of the brain vasculature, whereas spatial proximity, geneexpression similarity, and hemodynamic similarity do not explain the observed patterns (Fig. 2c). For breast cancer, we have *ρ*_distance_ = − 0.53 (*p*_moran_ = 0.94), *ρ*_FC_ = 0.48 (*p*_moran_ = 0.11), *ρ*_ATT_ = 0.58 (*p*_moran_ = 0.033), *ρ*_CBF_ = 0.51 (*p*_moran_ = 0.033), and *ρ*_gene_ = 0.49 (*p*_moran_ = 0.12). For lung cancer, we have *ρ*_distance_ = − 0.54 (*p*_moran_ = 1.0), *ρ*_FC_ = 0.45 (*p*_moran_ = 0.90), *ρ*_ATT_ = 0.61 (*p*_moran_ = 0.033), *ρ*_CBF_ = 0.59 (*p*_moran_ = 0.037), and *ρ*_gene_ = 0.53 (*p*_moran_ = 0.12). For melanoma, we have *ρ*_distance_ = − 0.40 (*p*_moran_ = 1.0), *ρ*_FC_ = 0.33 (*p*_moran_ = 0.99), *ρ*_ATT_ = 0.29 (*p*_moran_ = 0.94), *ρ*_CBF_ = 0.25 (*p*_moran_ = 0.90), and *ρ*_gene_ = 0.37 (*p*_moran_ = 0.90). All reported *p*-values are FDR-corrected. Together, the results show that the spatial organization of metastasis frequency maps for breast and lung cancers is conditioned by vascular network organization of the brain, such that regions with similar vascular properties display similar metastatic burden.

Although inter-regional covariance in gene transcription did not directly explain the spatial distribution of metastasis, this finding does not rule out a potential role for specific genes in shaping regional vulnerability to metastasis. Particular genes or biological pathways, rather than overall gene-expression similarity, may contribute to local tissue conditions that facilitate metastatic arrest, adhesion, or colonization. In Fig. S2 we therefore perform a more targeted gene enrichment analysis to examine whether spatial organization of metastasis frequency maps is associated with specific biological pathways, rather than genome-wide expression similarity.

All inter-regional similarity networks included in Fig. 2 comprised both cortical and cerebellar regions, given their involvement in metastatic brain disease. As a complementary analysis, we further perform a cortical-only node-neighbour analysis, allowing us to include additional similarity networks based on metabolic, receptor, and laminar annotations (Fig. S3). In this cortical-only analysis, CBF and ATT similarity networks continue to explain the breast and lung metastasis maps. Furthermore, metabolic similarity explains the observed cortical pattern of breast cancer metastasis, and receptor similarity explains the metastatic patterns in melanoma and lung cancer. Notably, after correcting for multiple comparisons across this broader set of cortical-only networks, none of the associations remains significant after FDR correction.

### Metastasis frequency maps and arterial border-zones

In the previous section, we showed that regional vulnerability to metastasis relates to the organization of the cerebral vasculature, which is spatially heterogeneous across the brain. Of particular interest are territories perfused by small-caliber vessels operating under relatively low hemodynamic pressures because they are more vulnerable to infiltration by emboli^13,126,127^. These regions, commonly referred to as arterial border-zones or watershed areas, occupy the interfaces between major vascular territories. In this section, we test whether metastatic lesions preferentially localize to the border-zones.

To quantify the spatial correspondence between metastasis frequency maps and arterial border-zones, we leverage arterial spin labeling (ASL) data from the HCP to derive *in vivo* estimates of arterial transit time (ATT). Arterial border-zones are delineated as regions exhibiting delayed arterial arrival times, reflecting the distal propagation of blood from the carotid arteries^32^. A normative ATT map is obtained by applying principal component analysis to concatenated individual-level maps (*N* = 597) and retaining the first principal component (Fig. 3a, see *Methods*). Metastasis frequency maps show a robust correspondence with the ATT map across both cortical and cerebellar regions for breast and lung cancers (breast: *ρ* = 0.50, *p*_moran_ = 5.49 × 10^−3^; lung: *ρ* = 0.59, *p*_moran_ = 5.49 × 10^−3^; melanoma: *ρ* = 0.30, *p*_moran_ = 0.11; all reported *p*-values are FDR-corrected) (Fig. 3b-e). These associations exceed those expected under spatial autocorrelation-preserving null models. In Fig. S5, we further incorporate a normative CBF map and its interaction with ATT, showing that neither CBF nor the ATT × CBF term exhibits a significant relationship with metastatic frequency. Taken together, these findings point to a central role for vascular architecture in shaping metastatic vulnerability. Specifically, arterial border-zones—characterized by distal perfusion and smaller-caliber vessels—may create a hemodynamic environment that favors the arrest and extravasation of circulating tumor cells, especially in breast and lung cancers. The physical organization of the arterial vasculature, rather than the magnitude of blood flow, is the dominant factor shaping metastatic frequency patterns in these cancers.

**Figure 3.**
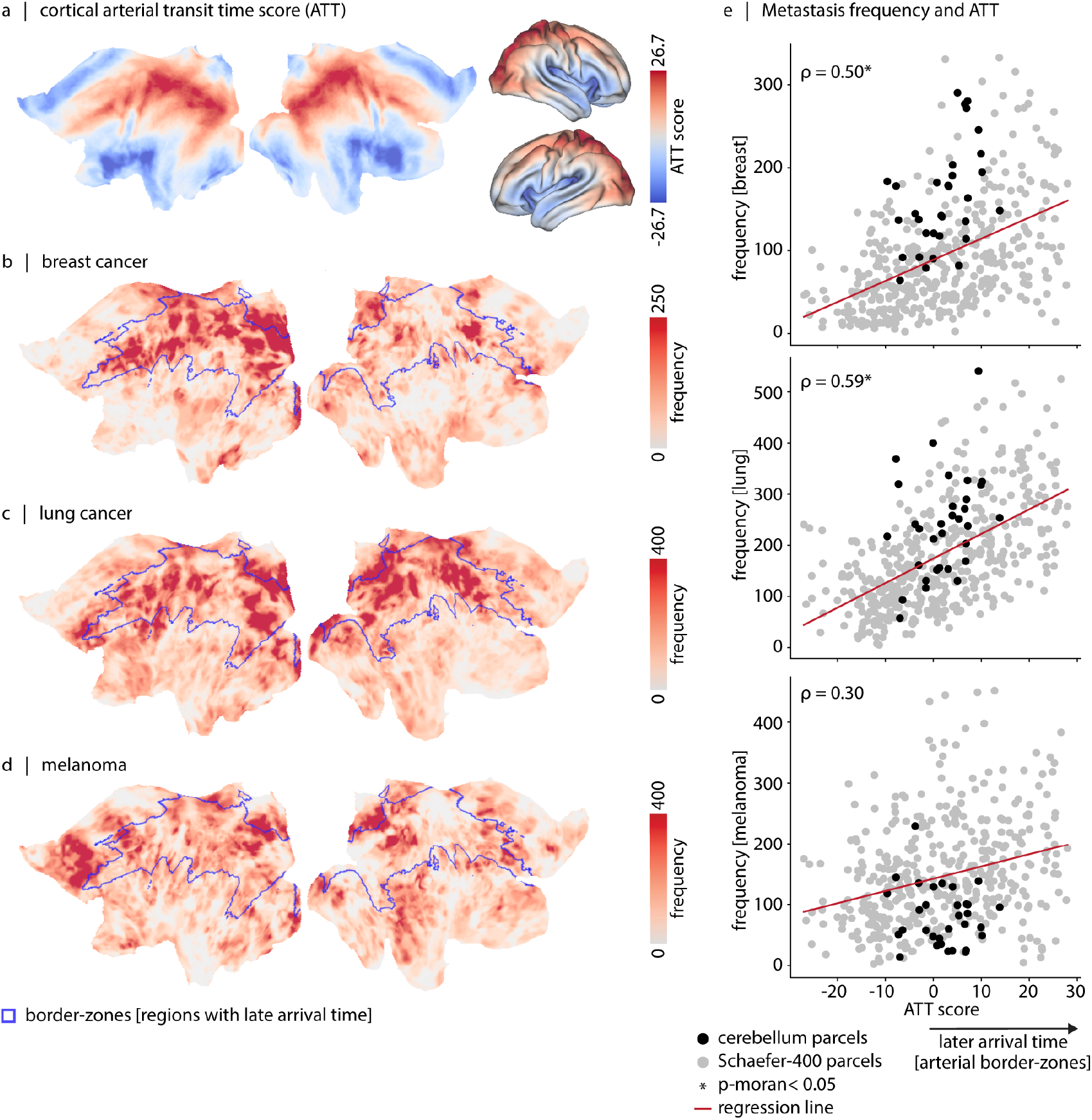
Brain metastasis frequency in primary breast and lung cancers correlates with arterial border-zones. (a) Arterial transit time (ATT) map displayed on 2D flat and 3D midthickness fsLR cortical surfaces. Brain metastasis maps for (b) breast cancer, (c) lung cancer, and (d) melanoma. Blue outlines in the ATT map highlight regions with longer arterial transit time. (e) Each scatter plot shows the Spearman’s rank correlation (*ρ*) between brain metastasis frequency maps (*y*-axis) and arterial transit time (*x*-axis). Each dot represents a brain region (gray: cortical Schaefer-400 parcels; black: 32 cerebellar parcels). Asterisks (*) indicate statistical significance based on spatial autocorrelation-preserving null models (FDR-corrected *p*_moran_ < 0.05). In Fig. S4, we show the cerebellar arterial transit time together with cerebellar metastasis frequency maps on cerebellar flat surfaces.

## DISCUSSION

Brain parenchymal metastasis is a frequent complication in patients with cancer, specifically those with primary tumors of the breast, lung, and melanoma^80,106^. Although primary tumors exhibit substantial molecular and genomic heterogeneity^7,67^,, metastatic spread to the brain shows consistent spatial patterns at the population level for each cancer type^6,8^. By aggregating data across a large cohort (*N*_*patients*_ > 2, 300), Barrios and colleagues estimated spatially heterogeneous maps of metastatic frequency across the brain for three primary cancer types, including breast, lung, and melanoma^8^. In this study, we sought to identify the biological features that render certain brain regions more susceptible to metastatic colonization. To this end, we examined multiple brain inter-regional similarity matrices, including transcriptomic similarity, hemodynamic similarity, and vascular architectural similarity, and tested whether regions that are biologically alike also share similar levels of metastatic burden. The spatial distribution of metastases is closely linked to the vascular architecture of the brain, particularly in primary breast and lung cancers. Notably, watershed regions, characterized by prolonged arterial transit times, exhibit the greatest frequency of metastatic lesions. These regions are supplied by small-caliber vessels and are located at the interface of major arterial territories.

Metastasis is a multi-step process whereby primary tumor cells disseminate through the blood or lymphatic circulation, arrest in distant organs, extravasate into the surrounding tissue, and ultimately survive and proliferate within a new microenvironment^18,34^ . Despite the systemic nature of dissemination, metastases do not occur uniformly across tissues but instead preferentially localize to specific organs, a phenomenon historically conceptualized as the “fertile soil” hypothesis. According to this framework, successful colonization depends on the compatibility between the intrinsic properties of disseminated tumor cells (the “seed”) and the local microenvironment of the host tissue (the “soil”)^22,87^. Importantly, this heterogeneity extends beyond organ-level selectivity to regional variation within individual organs. For example, in the liver, metastases preferentially localize to the right hepatic lobe^91,125^. A similar principle applies to brain metastasis, where different primary cancers give rise to distinct and spatially non-uniform patterns of lesion distribution (Fig. 1a)^8^. The genetic composition of the disseminated tumor cells and the biochemical environment of the brain are among the factors that contribute to this organ- and region-specific vulnerability. For example, tumor cells may be attracted to specific regions by chemo-attractants that facilitate transient attachments of tumor cells, their survival, and sustained growth. Beyond genetic and molecular determinants, mechanical and vascular factors also guide metastatic spread. According to the “mechanical hypothesis”^30^, disseminated tumor cells can become trapped within the microvasculature, with local hemodynamic conditions influencing their arrest and subsequent extravasation.

In the present study, we use a node-neighbour similarity framework to identify the biological substrates underlying regional vulnerability to brain metastasis. We construct inter-regional similarity networks where each edge encodes the pairwise similarity between two regions with respect to a given biological feature (such as gene expression or vascular features). By integrating inter-regional similarity networks with brain metastasis frequency maps^8^, we show that brain regions with similar vascular architecture have comparable levels of metastatic burden in primary breast and lung cancers, even after controlling for the background effect of spatial autocorrelation. The node-neighbour framework has previously been applied in studies of neurodegenerative diseases to assess whether biologically similar brain regions also exhibit similar levels of pathology^31,49,109,121^. Unlike conventional map-to-map analyses, which quantify spatial correspondence while treating regions as independent observations, the node-neighbour framework further incorporates inter-regional relational structure. It thereby captures how the biological neighbourhood of a region—defined by its similarity-based relationships to the rest of the brain—shapes its susceptibility to disease expression, or in the present context, to metastatic burden.

Brain metastasis frequency in primary breast and lung cancers co-localizes with arterial border-zones. This spatial pattern has been qualitatively described in the cancer literature^22,55^, but has not been systematically quantified at the whole-brain level, in part because *in vivo* imaging and localization of vascular territories and border-zones have been challenging due to the relatively low signal-to-noise ratio of arterial spin labeling (ASL) imaging. Here, we use high-quality HCP data from nearly 600 participants^14,62^, and construct a continuous, high-resolution normative map for arterial transit time. This map provides a hemodynamic quantity that directly indexes distance from major arterial supply and thereby captures the spatial organization of arterial border-zones^32^. We then quantify the spatial correspondence between metastasis frequency and arterial transit time across the whole brain, and show that regions with longer transit times exhibit higher metastatic burden. This relationship is consistent with vascular constraints on circulating tumor cell dynamics and arrest, as described by the “mechanical hypothesis”^30^. In large-caliber vessels, the rapid bloodstream reduces the likelihood of circulating tumor cell adhesion by effectively clearing them from the vasculature; however, arterial border-zones, supplied by small-diameter vessels with reduced flow, lack this mechanical protective effect^36^. The observation of further metastasis involvement in border-zone areas, complements the prior finding^8^ showing that brain metastatic foci are over-represented near the gray–white matter interface. This interface represents a vascular transition zone, where the vasculature becomes sparser as vessels enter white matter tissue and may therefore create favorable conditions for trapping circulating tumor cells^55,63^. Together, enrichment near arterial border-zones and the gray–white matter interface imply that the vascular organization of the brain, including vessel caliber, vascular density, and blood-flow dynamics, may shape the regional landscape of metastatic seeding. Notably, this association is not observed for melanoma, suggesting potential differences in the metastatic spread and mechanical trapping mechanisms across primary cancer types. Prior reports indicate that melanoma metastases show a stronger tendency to involve cortical and superficial regions^80^, consistently, high-resolution MRI work has shown that more than 90% of small intracranial melanoma metastases develop in close proximity to the leptomeninges, raising the possibility of a corticomeningeal route of dissemination^66^.

The spatial correspondence between arterial border-zones and regions of elevated metastatic frequency in breast and lung cancers points to a potential role of vascular physiology in shaping metastatic vulnerability. Specifically, it suggests that systemic vascular health and metabolic status of the body should be included in cancer metastasis studies as factors that may influence the likelihood of cell trapping. A plausible mechanistic basis for this link arises from the effects of metabolic dysfunction on cerebral hemodynamics and vascular integrity. In our previous work^32,33^, we showed that, beyond aging, elevated body mass index (BMI) and metabolic syndrome are associated with reduced cerebral blood flow and changes in the hydrodynamics of blood flow, particularly within watershed regions. Consistently, metabolic dysfunction has been shown to impair the neurovascular unit, promote blood–brain barrier breakdown, increase vessel permeability, and promote chronic inflammation in the brain^17,47,9^ —factors that may facilitate the arrest and extravasation of circulating tumor cells. The relationship between vascular comorbidities and brain metastatic potential in pathological conditions is not sufficiently characterized^76^. Nevertheless, emerging clinical evidence shows that obesity may increase the risk of distant metastasis in women with breast cancer^86,94^, further supporting a role for systemic metabolic factors in metastatic progression. Future work is needed to conclude whether the propensity for metastatic arrest within arterial border-zones is modulated by systemic metabolic health, and to identify peripheral biomarkers that may predict metastatic risk in individual patients.

In the present work, we found that brain regions with globally similar gene expression profiles do not necessarily show similar metastasis frequencies. Importantly, this result does not imply that genetic factors are completely unrelated to the spatial organization of metastasis. Rather, it suggests that large-scale inter-regional transcriptomic similarity may not directly account for these patterns. Analyses of correspondences between metastasis frequency maps and individual gene-expression maps revealed consistent enrichment of biological processes related to lipid metabolism, macrophage-like immune activity, and vascular biology in metastasis-vulnerable regions, particularly for breast and lung cancers. For melanoma, the prominence of PI3K-related processes among the top-ranked terms is consistent with the established involvement of this process in melanoma metastatic progression^21,61,^ . Overall, these results imply that metastatic patterns in the brain are shaped by an interplay between global vascular architecture and local tissue biology.

The present work should be considered alongside several methodological considerations. First, the interregional similarity matrices, normative arterial transit time map, and gene expression maps are estimated from independent cohorts of healthy individuals rather than from the patient populations used to derive the metastasis frequency maps. At present, acquiring and integrating multi-modal biological measurements within the same individuals remains challenging. The challenges originate from practical constraints in patient populations and from the fact that typical metastasis studies rarely incorporate advanced neuroimaging modalities, such as arterial spin labeling or functional MRI. The normative maps employed in this study may not fully capture the hemodynamic and transcriptomic landscape of the diseased brain. Second, the brain metastasis frequency maps presented here aggregate patients with heterogeneous immunohistochemical tumor profiles. Primary cancers are molecularly heterogeneous and comprise subtypes with distinct metastatic behaviors^64,115^. For example, among non–small cell and small cell lung cancer, the former has a high risk of spreading to the cerebellar hemisphere^40,106^. Collapsing across such subtypes may therefore obscure biologically meaningful variation in spatial patterns of metastasis and preclude assessment of inter-individual variability. Emerging open-access initiatives that integrate molecular, clinical, and imaging data within the same individuals, such as The Cancer Imaging Archive (TCIA^20^; https://www.cancerimagingarchive.net), will support a more precise characterization of subtype-specific and patient-level metastatic patterns. Last, throughout the paper, we used correlative methods to investigate the spatial associations between cancer metastasis frequency maps and biological brain features, and therefore cannot establish the causal mechanisms underlying these associations. Future experimental models of brain metastatic seeding (e.g., hemodynamic perturbation models)^36,90^ can help characterize the influence of vascular and genetic organization on the initial deposition of circulating tumor cells and their subsequent survival and post-deposition growth.

## METHODS

All code used to perform the analyses is available on GitHub at https://github.com/netneurolab/Farahani_Brain_Metastases.

### Brain metastasis frequency maps

Brain metastasis frequency maps for primary breast cancer, lung cancer, and melanoma were obtained from Barrios and colleagues and are publicly available at https://osf.io/fkqmr^8^. In short, the dataset included patients presenting for first-time stereotactic radiosurgery following a new diagnosis of brain metastasis at four academic medical centers in the United States and Canada between January 2011 and December 2023. Patients had a median of 2 metastases (range 1–53) with a median lesion volume of 0.23 cc (range 0.002–62.381). Breast patients had the highest mean= 5/median= 3 lesion count combination (range 1–53). Lesions were identified on pre-radiosurgery post-gadolinium contrast T1– weighted MRI and manually delineated by radiation oncologists. MR images were bias-corrected and intensity-normalized. Metastasis frequency maps were generated by aggregating lesion occurrences across all patients in standard space, yielding voxel-wise distributions of metastatic burden. Further details on cohort characteristics are provided in Table. S1, this table is derived from the original manuscript. Imaging protocols and map construction details are provided in the original study^8^.

### HCP–A: Demographics

We analyzed data from 597 participants (329 females; 268 males) aged 36–100 years from the HCP–Aging dataset (HCP Lifespan studies, 2.0 Release)^14,50,^ . Participants represented a “typical” aging population and had common health conditions (e.g., hypertension) without identified pathological causes of cognitive decline (e.g., stroke). All study procedures were conducted in accordance with the principles expressed in the Declaration of Helsinki and were approved by the Institutional Review Board at Washington University in St. Louis.

### HCP–A: Brain imaging acquisition

All HCP–A brain imaging data were acquired using a 3.0 Tesla Prisma scanner (Siemens; Erlangen, Germany) and a 32–channel Prisma head coil^14,50,^ .

T1–weighted structural data were acquired using a multi-echo magnetization-prepared rapid gradient echo (MPRAGE) sequence with the following parameters: repetition time (TR) = 2 500 ms, inversion time (TI) = 1 000 ms, echo times (TE) = 1.8/3.6/5.4/7.2 ms, spatial resolution = 0.8 × 0.8 × 0.8 mm^3^, number of echoes = 4 and flip angle = 8^°^. T2–weighted structural data were acquired using a 3D sampling perfection with application-optimized contrasts using different flip angle evolutions (SPACE) sequence, with the same spatial resolution as the T1–weighted image. Parameters for the T2–weighted sequence were: TR = 3 200 ms, TE = 564 ms and turbo factor = 314. Both T1– and T2–weighted images captured a sagittal field of view measuring 256 × 240 × 166 mm, and a matrix size of 320 × 300 × 208 slices. Additional acquisition parameters included 7.7% slice over-sampling, 2–fold in-plane acceleration (GRAPPA) in the phase encoding direction, and a 744 Hz/Px pixel bandwidth. T1– and T2–weighted structural data provided the anatomical reference for analysis of all imaging modalities, reconstruction of cortical surfaces, and estimation of cortical myelin content (T1/T2 ratio) and cortical thickness^50^.

Resting-state fMRI data with blood-oxygen-level-dependent (BOLD) contrast were acquired using a 2D multi-band gradient-recalled echo (GRE) echo-planar imaging (EPI) with the following parameters: TR/TE = 800/37 ms, flip angle = 52^°^, spatial resolution = 2.0 × 2.0 × 2.0 mm^3^. Functional scans were acquired in pairs of two runs (four runs in total per participant, each run lasting 6.5 min), with opposite phase encoding polarity so that the fMRI data in aggregate were not biased toward a particular phase encoding polarity (two runs had the phase encoding of anterior-to-posterior (AP) and two runs had the phase encoding of posterior-to-anterior (PA)). During rs-fMRI scanning, participants viewed a small white fixation crosshair on a black background. The functional MRI minimal preprocessing steps that were applied to the data are provided by Glasser and colleagues^43^.

Blood perfusion and arterial transit time were measured using arterial spin labeling (ASL) magnetic resonance imaging (MRI)^58,62^. ASL data were acquired using a pseudo-continuous arterial spin labeling (pCASL) and 2D multi-band (MB) echo-planar imaging (EPI) sequence. Pseudo-continuous ASL data were acquired with labeling duration of 1 500 ms and five post-labeling delays of 200 ms, 700 ms, 1 200 ms, 1 700 ms, and 2 200 ms, containing 6, 6, 6, 10, and 15 control-label image pairs, respectively. To calibrate perfusion measurements into units of ml/100g/min, two PD–weighted M0 calibration images (TR > 8 s) were acquired at the end of the pCASL scan. Other sequence parameters included: spatial resolution = 2.5 × 2.5 × 2.5 mm^3^, and TR/TE = 3 580/18.7 ms. For susceptibility distortion correction, two phase-encoding-reversed spin-echo images were also acquired. Participants viewed a small white fixation crosshair on a black background during the scan time (5.5 min). The ASL data preprocessing was conducted following the ASL Pipeline for the Human Connectome Project available at https://github.com/physimals/HCP-asl (explained in detail in Kirk and colleagues^62^). To run this pipeline, we used QuNex platform (singularity container, version 0.99.1)^57^.

### Distance matrix

The distance matrix quantifies the Euclidean distance between pairs of brain parcel centroids, where higher values indicate greater spatial separation. Parcel centroids were estimated as the center of mass of each parcel in the combined cortical and cerebellar atlas in MNI volumetric space, computed using the center_of_mass function from scipy.ndimage. Pairwise Euclidean distances between all parcel centroid coordinates were then calculated using the cdist function from `scipy.spatial.distance`. The diagonal was set to zero. The resulting matrix is shown in Fig. 2a (leftmost heatmap).

### Hemodynamic similarity

Hemodynamic similarity, often referred to as functional connectivity, quantifies the similarity between brain regions based on the synchronization of their BOLD signal fluctuations. Resting-state functional MRI data were obtained from the HCP–A dataset. Vertex-wise time-series were parcellated into regional time-series and demeaned per imaging run (each subject underwent four runs). The regional time-series were then concatenated across runs and *z*-scored for each participant. Each participant’s unified time-series was used to estimate a functional connectivity matrix. Functional connectivity matrices were computed for each participant using the Pearson correlation between all pairs of regional time-series. A group-level connectivity matrix was then obtained by averaging the functional connectomes across participants. The resulting group-level matrix is shown in Fig. 2a (second heatmap from the left).

### Arterial transit time and cerebral blood flow similarity

ATT and CBF similarity matrices quantify vascular similarity between pairs of brain regions. ATT and CBF maps were obtained from the HCP–A dataset. These matrices were constructed by calculating the similarity of regional ATT or CBF profiles across participants, such that regions exhibiting similar inter-individual variation in ATT/CBF were considered more strongly connected.

Specifically, to construct these matrices, vertex-wise ATT/CBF data were first parcellated by averaging signals within each parcel of the combined cortical and cerebellar atlas, yielding a parcel × participant data matrix. To account for inter-individual differences in global ATT/CBF levels, regional profiles were *z*-scored across participants (i.e., within each parcel), such that each region’s profile reflected relative rather than absolute ATT/CBF values. The resulting normalized matrix was then used to compute a parcels × parcels covariance matrix, where each entry encodes the covariance between the cross-participant perfusion profiles of two regions. The underlying premise is that regions characterized by common arterial properties tend to exhibit correlated inter-individual variation in CBF and ATT, such that the resulting similarity matrix captures shared vascular supply architecture. Consistent with this interpretation, we have previously shown that gradients estimated from the CBF similarity network recapitulate known arterial territories^32^. The resulting matrices are shown in Fig. 2a (second and third matrices from the right).

### Gene expression similarity

Gene expression similarity matrix quantifies the transcriptomic similarity between pairs of brain regions. The underlying data were obtained from bulk tissue microarray expression data collected from six post-mortem brains (1 female; age: 24–57, mean age: 42.50±13.38 years). Data were provided by the Allen Human Brain Atlas (https://human.brain-map.org)^52^ and were processed using the abagen toolbox, available at https://github.com/rmarkello/abagen^71^, yielding a map for each gene in the parcellated MNI template. Only genes with high differential stability across donors (threshold of 0.1) were retained, resulting in 12, 803 genes. Similarity between regions was computed using the Pearson correlation coefficient between their gene expression profiles, producing a region × region matrix. The resulting matrix is shown in Fig. 2a (rightmost heatmap).

### Metabolic similarity

Metabolic similarity quantifies the similarity between cortical brain regions based on their glucose metabolism. Data were obtained from [^18^F]-fluorodeoxyglucose positron emission tomography (PET) imaging. The dataset included 26 healthy participants (77% female; age: 18–23) who participated in a 95-minute simultaneous MR-PET scan acquired using a 3T molecular MR scanner^56^. PET images were preprocessed according to previous work^122^. Each volume of the PET time-series was registered to the MNI152 template space and was parcellated according to the Schaefer-400 atlas^105^. Regional time-series were then correlated using the Pearson correlation to construct a metabolic connectivity matrix for each participant. A group-level metabolic connectivity matrix was then obtained by averaging the metabolic connectomes across participants^49^. The resulting matrix is shown in Fig. S3a (rightmost heatmap).

### Receptor similarity

Receptor similarity quantifies similarity between cortical brain regions based on neurotransmitter receptor density profiles. To construct this network, PET tracer images for 18 neurotransmitter receptors and transporters were used^48^. These receptors/transporters covered nine neurotransmitter systems, including dopamine (D1, D2, DAT), norepinephrine (NET), serotonin (5-HT1A, 5-HT1B, 5-HT2, 5-HT4, 5-HT6, 5-HTT), acetylcholine (*α*4*β*2, M1, VAChT), glutamate (mGluR5), GABA (GABAA), histamine (H3), cannabinoid (CB1), and mu-opioid (MOR). See Table. S2 for a list of the receptors and transporters used. Each of these PET tracer images was parcellated based on the Schaefer-400 atlas^105^ and normalized using *z*-scores. Similarity between regions was computed using the Pearson correlation between receptor profiles, yielding a region × region matrix^49^. The resulting matrix is shown in Fig. S3a (second matrix from the right).

### Laminar similarity

Laminar similarity quantifies similarity between cortical brain regions based on their cellular distributions across cortical layers^49,89^. Data came from the high-resolution (20 µm) histological BigBrain atlas, a postmortem Merker-stained histological atlas of a 65-year-old male^3^. Staining intensity profiles were sampled across 50 equi-volumetric surfaces within the cortical grey matter, enabling the assessment of neuronal density and soma size variations across cortical layers. These intensity profiles also helped delineate boundaries among cortical layers, such as supragranular (layers I–III), granular (layer IV), and infragranular (layers V–VI). The BigBrainWarp toolbox^88^ was used to transform the data to the surface-based fs-LR template, which was then parcellated based on the Schaefer-400 atlas^105^. A laminar similarity matrix was estimated by computing the partial correlation between regional intensity profiles of pairs of cortical regions^49^. The resulting matrix is shown in Fig. S3a (third matrix from the right).

### Node-neighbour framework

The metastatic burden of a brain region (i.e., node) was defined as the mean metastasis frequency value within that region and was denoted as *d*_i_ for region_i_. For each similarity network (Euclidean distance, functional connectivity, ATT, CBF, and gene expression), negative edge weights were set to zero prior to analysis, retaining only positive connections. For region_i_, region_j_ was considered a neighbour if the pair had a positive connection strength defined based on the inter-regional similarity network. The metastatic exposure of region_i_ was computed as the weighted average metastatic burden of its neighbouring regions, with weights given by the strength of inter-regional similarity (*c*_ij_):

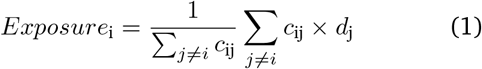

where *c*_ij_ represents the similarity between region_i_ and region_j_, and *d*_j_ is the metastatic burden of region_j_. To assess whether metastatic burden was organized according to the underlying similarity network, we computed the Spearman’s rank correlation between each region’s burden and its metastatic exposure (Fig. 2b):

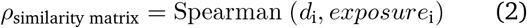

Statistical significance was assessed by comparing the empirical correlation against a null distribution of correlations derived from spatially constrained null metastasis maps. For each null map, the node-neighbour exposure was recomputed using the same similarity matrix, and the Spearman correlation between the null map and its exposure was recorded. The empirical *ρ* was then compared against this null distribution to obtain a *p*-value. Multiple comparisons across the 15 tests (five similarity matrices × three cancer types) were controlled using false discovery rate correction via the Benjamini– Yekutieli procedure.

### Arterial transit time and cerebral blood flow maps

To estimate a normative arterial transit time (ATT) map, individual vertex- and voxel-wise ATT maps were first *z*-scored and then concatenated into a single data matrix. Principal component analysis (PCA) was then applied to this matrix. The first principal component accounted for % of the variance in ATT maps across the HCP-A participants (the second component accounts for 3.57% of the variance). The first principal component score map is shown in Fig. 3a.

To estimate a common cerebral blood flow map, which is representative of all participants in the sample, we *z*-scored blood perfusion maps of individual participants and concatenated them into a single data matrix. Participant-level standardization mitigates perfusion-level offsets arising from biological sex differences. We next applied PCA dimensionality reduction to the data matrix. The first principal component accounted for 47.68% of the variance in cerebral blood flow across the sample (the second component accounted for 2.55% of the variance). The first component reflected the main regional pattern of blood flow shared across participants and was used in subsequent analyses illustrated in Fig. S5.

### Gene enrichment analysis

We identified biological pathways associated with brain metastasis frequency maps using gene category enrichment analysis (GCEA). Frequency maps and gene expression data were parcellated across both cortical and cerebellar regions, and genes with differential stability below 0.1 were excluded to ensure robustness. Gene categories were defined using established pathway annotations (“GCEA_GO-biologicalProcessProp-discrete”) (restricted to categories with at least five genes; *N*_pathways_ = 5, 846)^15^. For each pathway, we computed a category score by correlating the expression profiles of its constituent genes with the metastasis frequency map (using Spearman’s rank correlation, *ρ*). To assess statistical significance while accounting for spatial autocorrelation, we generated 10, 000 Moran spectral randomizations of the frequency maps. For each null map, we recomputed the category scores to construct a null distribution. Empirical category scores were then compared against this null distribution, and two-sided *p*-values were obtained. Multiple comparisons were controlled using the False Discovery Rate (FDR). This ensemble-based null model provides a more conservative and spatially informed assessment compared with approaches that rely on random gene sampling^37^. Specifically, this framework tests whether genes within a given category show stronger associations with the observed brain phenotype than expected under spatially constrained random phenotypes.

### Atlases

Cortical data were parcellated using the Schaefer-400 atlas^105^. Cerebellar data were parcellated using the probabilistic atlas of the cerebellar lobules introduced by Diedrichsen and colleagues^24^. Note that the cerebellar atlas comprises 34 regions; however, only 32 regions were included in analyses where statistical significance was assessed. This exclusion was necessary because implementing Moran spectral randomization required resampling the MNI template to a lower resolution to make distance calculations computationally feasible. The two smallest regions, the left and right fastigial nuclei, were removed at this step due to their small size (fewer than 6 voxels in the original 1 mm^3^ resolution atlas).

### Statistics and null models

Throughout this paper, we used Moran spectral randomization to generate random null maps^26,123^. The null maps were generated using the `neuromaps` toolbox (available at https://github.com/netneurolab/neuromaps)^72^. Moran spectral randomization is a parameterized method that relies on a spatially-informed weight matrix, typically defined as the inverse distance between brain regions. However, rather than estimating parameters via the least-squares approach, this method performs an eigen decomposition of the spatial weight matrix to derive spatial eigenvectors that estimate the underlying autocorrelation structure of the brain map of interest. These eigenvectors are then used to impose a similar spatial structure onto randomly generated surrogate data.

Throughout the paper, statistical significance was defined at *p*-value< 0.05. Multiple comparisons were controlled using false discovery rate (FDR) correction via the Benjamini–Yekutieli procedure, implemented in the `multipletests` function of the `statsmodels` toolbox (fdr-by option).

## Data and code availability

The Human Connectome Project-Aging (HCP–A) is accessible through https://www.humanconnectome.org/study/hcp-lifespan-aging^14,50,113^.

Parcellation files, including the Schaefer-400 atlas^105^ and the cerebellar lobular atlas^24^, are publicly available at https://github.com/ThomasYeoLab/CBIG/tree/master/stable_projects/brain_parcellation/Schaefer2018_LocalGlobal/Parcellations and https://github.com/DiedrichsenLab/cerebellar_atlases/tree/master/Diedrichsen_2009, respectively.

All code used to perform the analyses can be found on GitHub at https://github.com/netneurolab/Farahani_Brain_Metastases. The code directly relies on open-source Python packages including NumPy (version 1.21.6)^51,117^, SciPy (version 1.7.3)^120^, pandas (version 1.3.5)^75^, seaborn (version 0.12.2)^124^, Matplotlib (version 3.5.3)^54^, statsmodels (version 0.13.5)^108^, bctpy (version 0.6.1)^99^, Nilearn (version 0.10.1, see^1^), NiBabel (version 4.0.2)^16^, and netneurotools (version 0.2.3)^68^. All brain plots in the manuscript are visualized using Connectome Workbench (version 1.5.0), available at https://www.humanconnectome.org/software/get-connectome-workbench/^70^.

## Acknowledgments

We thank Filip Milisav, Eric G. Ceballos, Yigu Zhou, Tahmineh Taheri, and Moohebat Pourmajidian for their comments and suggestions on the manuscript. AF acknowledges support from the Harold and Audrey Fisher Brain Tumor Student Award. BM acknowledges support from the Natural Sciences and Engineering Research Council of Canada (RGPIN−2017 − 04265), Canadian Institutes of Health Research (PJT−180439), and Canada Research Chairs Program (CRC−2022 − 00169).

---

**Figure S1.**
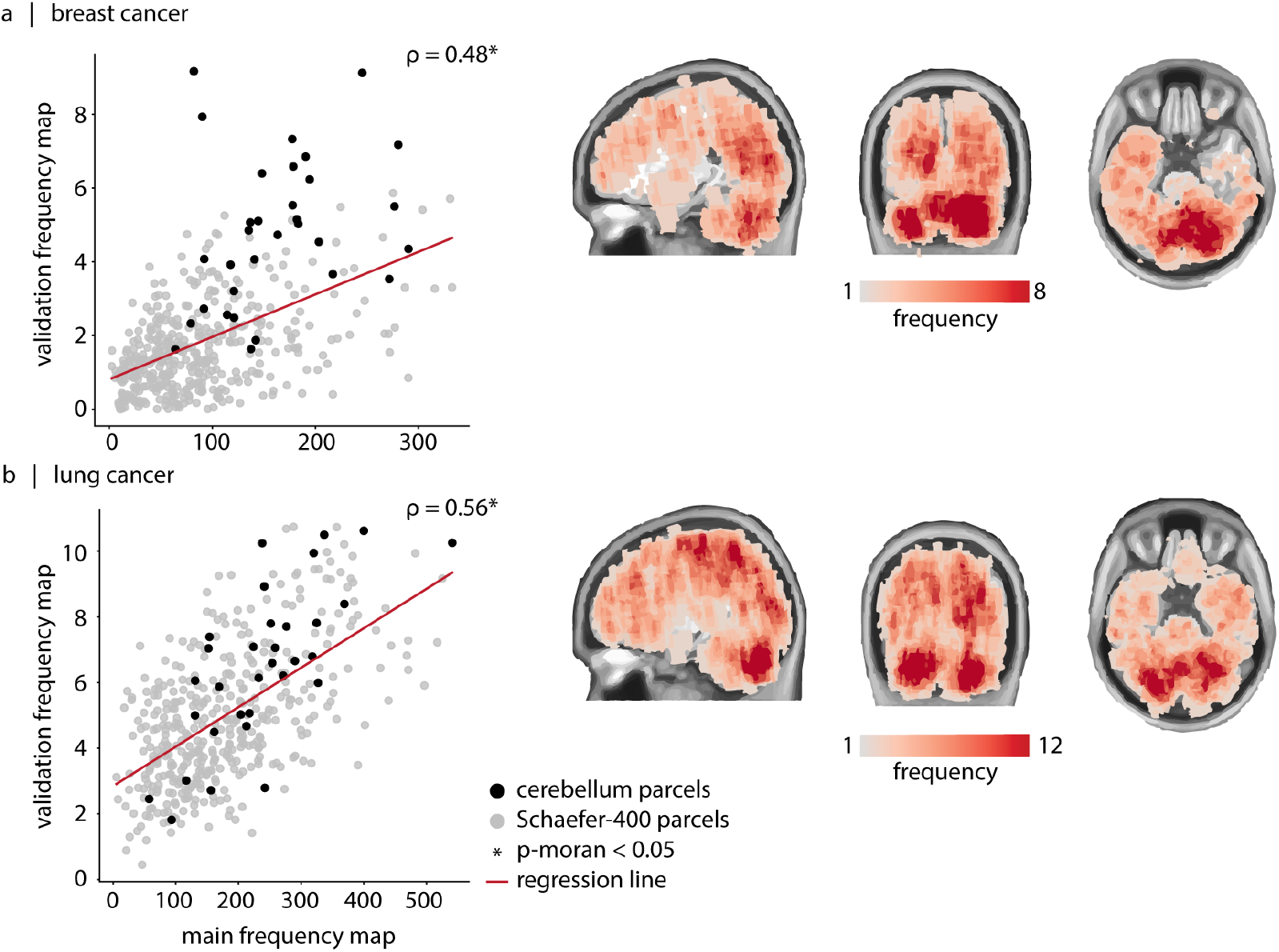
Cross-dataset correspondence of breast and lung brain metastasis frequency maps. The scatter plots show the correspondence between the metastasis frequency maps reported by Barrios and colleagues^8^ (*x*-axis) and those reported by Bao and colleagues^6^ (*y*-axis) (gray: cortical Schaefer-400 parcels; black: 32 cerebellar parcels). (a) The correlation between the two for primary breast cancer is equal to 0.48, and (b) the correlation between the two for primary lung cancer is equal to 0.56. Asterisks (*) indicate statistical significance based on spatial autocorrelation-preserving null models (*p*_moran_ < 0.05). Validation metastasis frequency maps from Bao and colleagues^6^ are shown on a T1-weighted group average template (MNI152; right panels in a and b).

**Figure S2.**
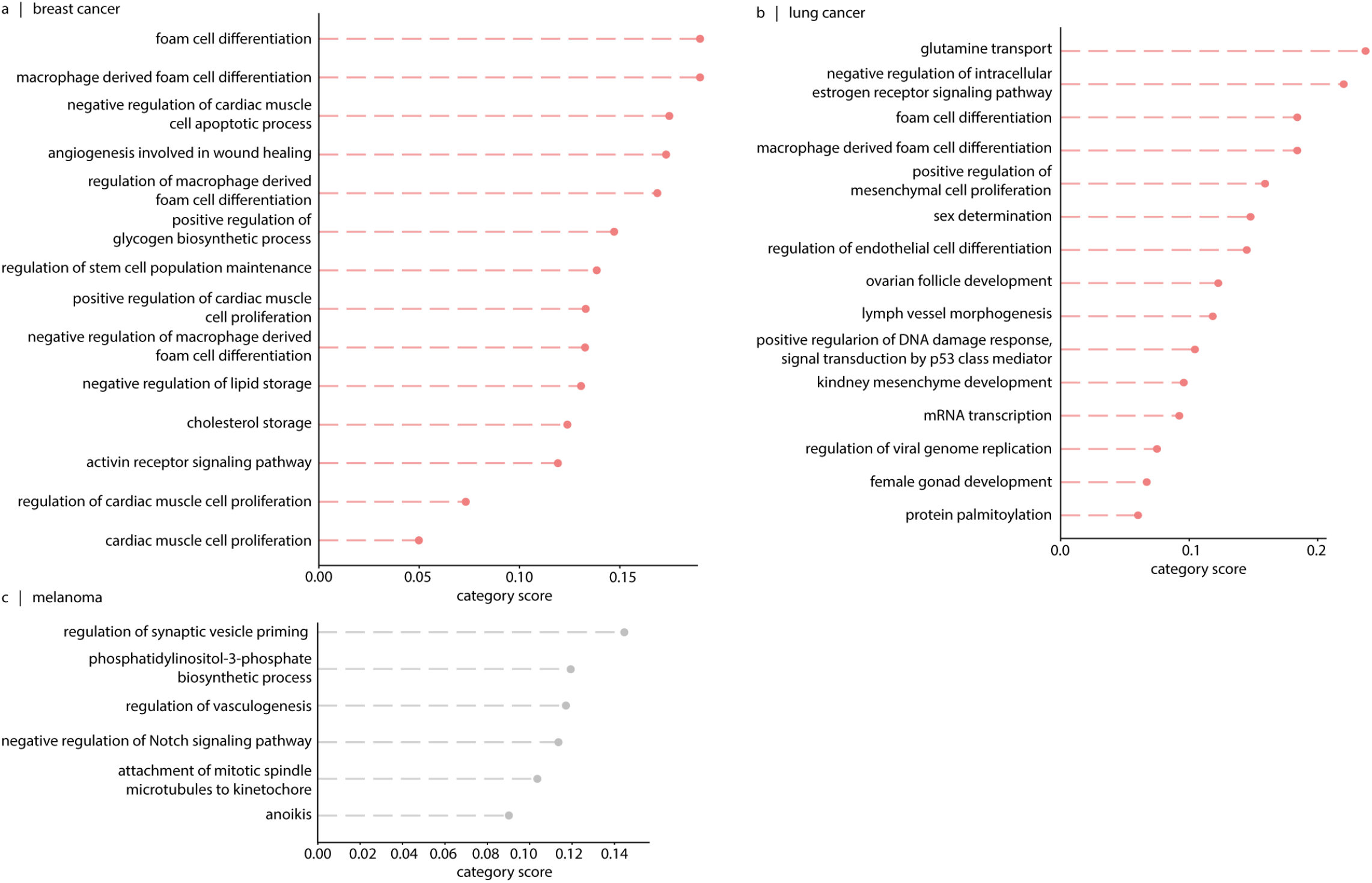
Biological pathways associated with brain metastasis frequency maps. We investigate the biological pathways associated with genes that are preferentially co-expressed in brain regions vulnerable to metastasis burden. Using pathway annotations provided in^15^, we identify biological pathways whose constituent genes show stronger correlations with the empirical metastasis maps than expected under spatial autocorrelation-preserving null models. The resulting *p*-values are further controlled for multiple comparisons using FDR correction. We find 14 and 15 significant biological pathways that co-localize with breast and lung metastasis maps (shown in orange), respectively. For breast cancer, the top-ranked pathways include “foam cell differentiation”, “macrophage derived foam cell differentiation”, and related regulatory pathways. Foam cells are lipid-laden macrophages arising from dysregulated intracellular lipid metabolism. These cells participate in impairing macrophage immune function and perpetuating inflammation^45^. Enrichment of these pathways may reflect involvement of lipid metabolism and macrophage-like immune pathways in shaping metastatic burden. Additional significant pathways are linked to vascular biology and lipid metabolism. Together, these findings point to the potential role of the immune system, lipid metabolism, and vascular features in shaping vulnerability to metastasis. For lung cancer, the top-ranked pathways again relate to foam cells. Furthermore, sex-related biological pathways, including “sex determination”, “negative regulation of intracellular estrogen receptor signaling pathway”, are among the top-ranked pathways. This observation may relate to the reported sex differences in brain metastatic rates of lung cancer, with women showing a higher proportion of brain metastases in non-small cell lung cancer^44^. We also find pathways that relate to the vascular system of the brain (i.e., “regulation of endothelial cell differentiation” and “lymph vessel morphogenesis”). For melanoma, no pathway reaches statistical significance after multiple-comparisons correction; nevertheless, the top-ranked processes can still provide biologically relevant insights and are reported here (shown in gray). Interestingly, “phosphatidylinositol-3-phosphate biosynthetic pathway (PI3K-related)” is among the top-ranked processes. This is consistent with prior evidence that shows associations between the phosphatidylinositol 3-kinase (PI3K) pathway and survival and proliferation of metastatic melanoma cells^21,61,98,116^, as well as with studies showing that targeting the PI3K/Protein Kinase B (AKT) pathway offers a therapeutic opportunity for patients with brain metastasis^19,83^.

**Figure S3.**
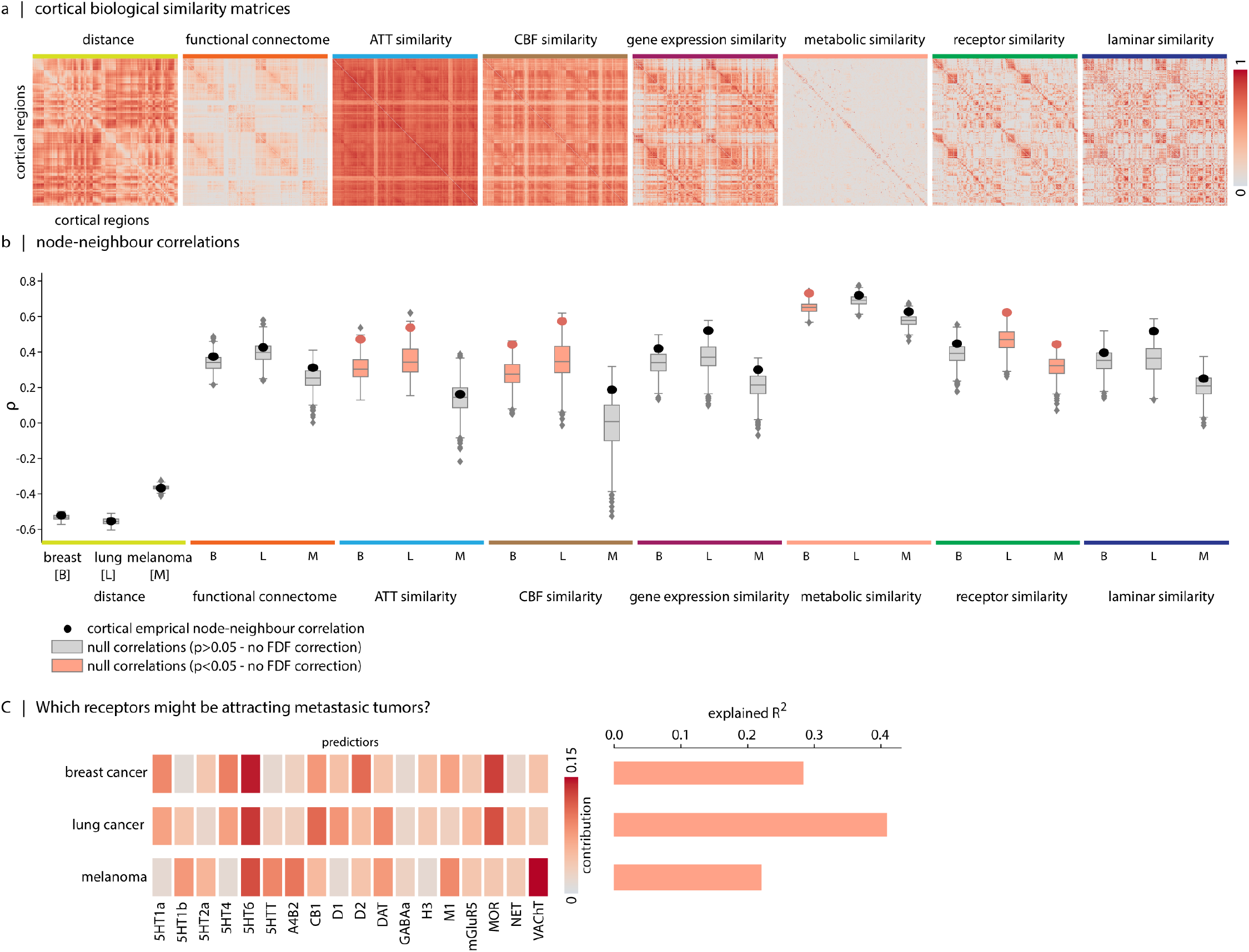
Do cortical biological similarity matrices explain the cortical metastasis frequency maps? We examine how brain cortical inter-regional biological similarity matrices shape the spatial organization of brain metastasis frequency maps. (a) Heatmaps of cortical biological similarity matrices, including Euclidean distance, hemodynamic, arterial transit time, cerebral blood flow, transcriptomic, metabolic, receptor, and laminar similarity. (b) Empirical correlations (*ρ*) between regional metastatic burden and neighbour-weighted burden, shown alongside null distribution of correlations. Boxplots in orange indicate statistical significance based on spatial autocorrelation-preserving null models (not FDR-corrected, *p*_moran_ < 0.05). For breast cancer, *ρ*_distance_ = − 0.52 (*p*_moran_ = 0.52), *ρ*_FC_ = 0.37 (*p*_moran_ = 0.45), *ρ*_ATT_ = 0.47 (*p*_moran_ = 0.009), *ρ*_CBF_ = 0.44 (*p*_moran_ = 0.023), *ρ*_gene_ = 0.42 (*p*_moran_ = 0.23), *ρ*_met_ = 0.73 (*p*_moran_ = 0.005), *ρ*_recep_ = 0.45 (*p*_moran_ = 0.32), and *ρ*_laminar_ = 0.40 (*p*_moran_ = 0.47). For lung cancer, *ρ*_distance_ = − 0.55 (*p*_moran_ = 0.92), *ρ*_FC_ = 0.43 (*p*_moran_ = 0.61), *ρ*_ATT_ = 0.54 (*p*_moran_ = 0.026), *ρ*_CBF_ = 0.57 (*p*_moran_ = 0.046), *ρ*_gene_ = 0.52 (*p*_moran_ = 0.052), *ρ*_met_ = 0.72 (*p*_moran_ = 0.31), *ρ*_recep_ = 0.62 (*p*_moran_ = 0.017), and *ρ*_laminar_ = 0.52 (*p*_moran_ = 0.07). For melanoma, *ρ*_distance_ = − 0.37 (*p*_moran_ = 0.77), *ρ*_FC_ = 0.31 (*p*_moran_ = 0.32), *ρ*_ATT_ = 0.16 (*p*_moran_ = 0.78), *ρ*_CBF_ = 0.19 (*p*_moran_ = 0.18), *ρ*_gene_ = 0.30 (*p*_moran_ = 0.24), *ρ*_met_ = 0.63 (*p*_moran_ = 0.12), *ρ*_recep_ = 0.44 (*p*_moran_ = 0.049), and *ρ*_laminar_ = 0.25 (*p*_moran_ = 0.52). All reported *p*-values are uncorrected. After FDR correction for multiple comparisons, none of the associations remained significant. (c) Given the explanatory role of receptor similarity shown in (b), we next aim to identify receptor maps that may contribute to the spatial patterning of metastasis maps. We fit multi-linear regression models predicting metastasis tumor distributions from receptor distributions. The significance of the models was assessed using spatially constrained null models, with all three models reaching statistical significance. Dominance analysis is then used to partition the model fit across predictors (receptor maps), enabling comparison of their relative contributions. The percent contribution of each predictor is defined as its normalized dominance relative to the total model fit.

**Figure S4.**
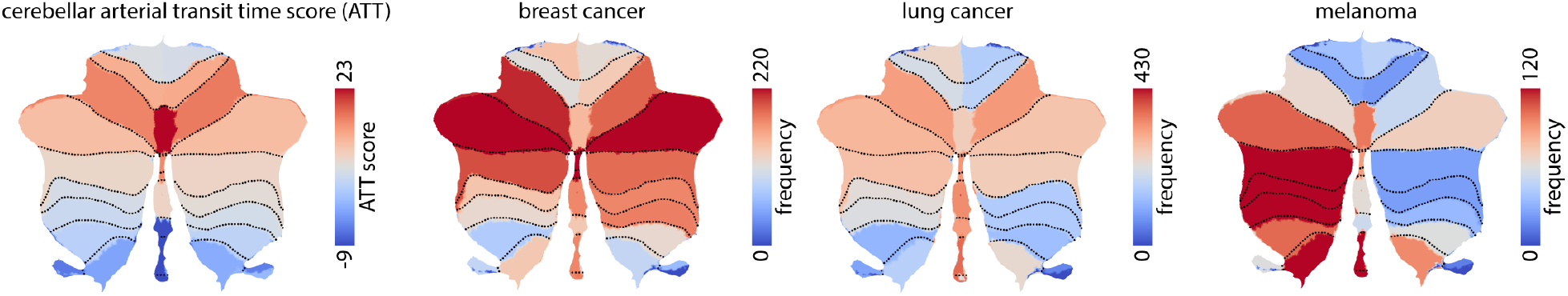
Cerebellar metastasis frequency and arterial border-zones. ATT score map is shown in the leftmost panel. Regions in red represent regions with delayed blood arrival. Cerebellar metastasis frequency maps for breast cancer, lung cancer, and melanoma are shown on the cerebellar flat surfaces.

**Figure S5.**
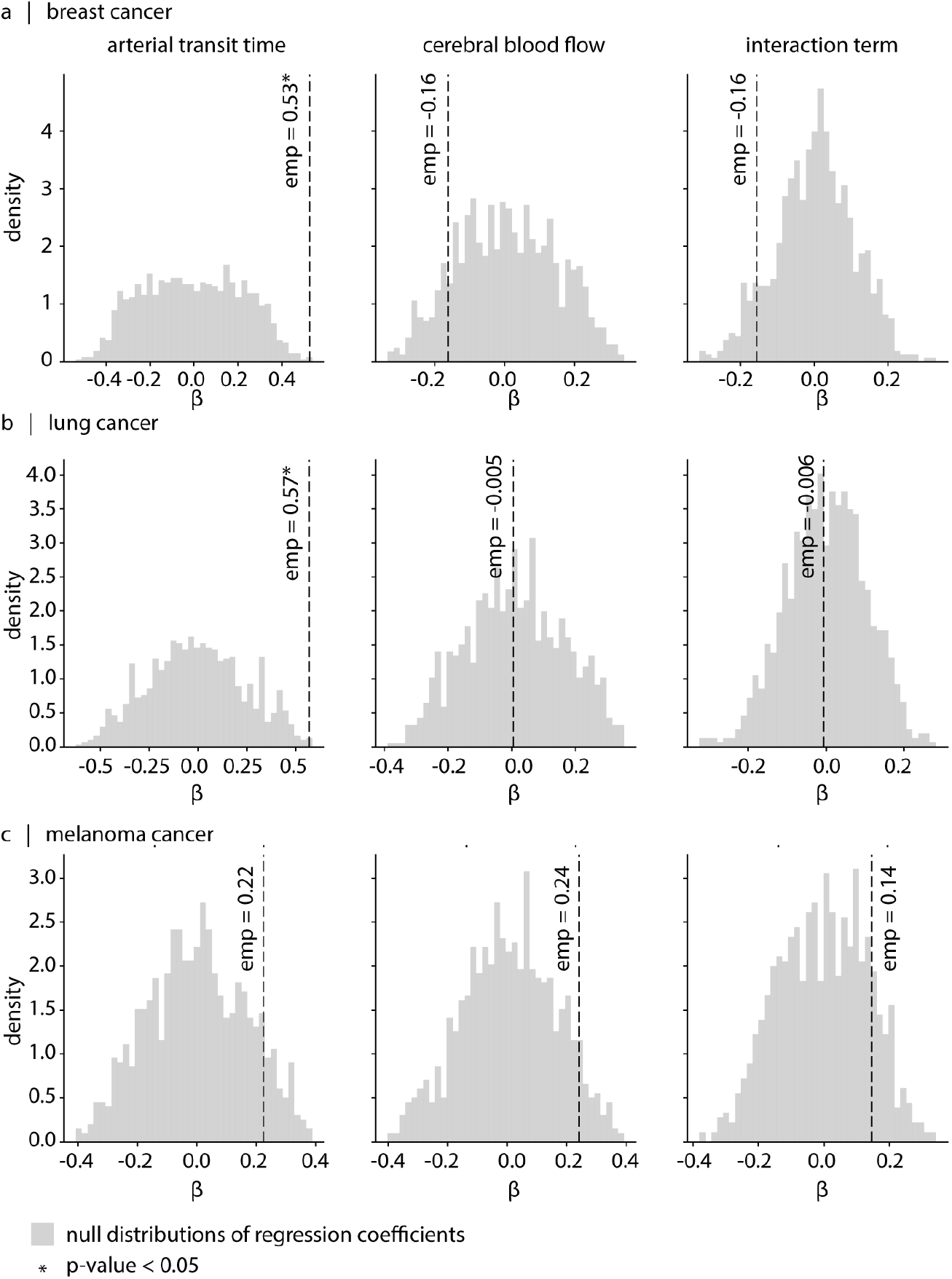
Contribution of arterial transit time, cerebral blood flow and their interaction in predicting the brain metastasis frequency maps. We constructed a linear model of the form: Cancer frequency = *β*_0_ +*β*_ATT_ ATT+*β*_CBF_ CBF+*β*_CBF×ATT_ (CBF × ATT). We evaluated the statistical significance of model coefficients by comparing the empirical estimates against a null distribution generated using Moran spectral randomization (*N*_moran_ = 1, 000). Specifically, the cancer frequency map was replaced with spatially autocorrelation-preserving null maps, and the model was recomputed for each iteration. The results indicate that the coefficient associated with arterial transit time (*β*_ATT_) is significantly greater than expected under the null for both breast and lung cancers.

**TABLE S1.** Cohort characteristics across sites and cancer types. See the original publication for further details^8^.

| Characteristic | MSKCC (N=371) | PMH (N=373) | UCD (N=377) | UCSF (N=1,184) | Total (N=2,305) |
| --- | --- | --- | --- | --- | --- |
| **Breast** |  |  |  |  |  |
| Patients, N | 87 | 64 | 74 | 348 | 573 |
| Lesions, N | 382 | 309 | 267 | 2,120 | 3,078 |
| Age, mean (range) | 54 (30–81) (N=81) | 58 (33–83) (N=40) | 56 (30–87) | 53 (0–79) | 54 (30–87) (N=543) |
| Sex, N (M/F) | 0/87 | 1/63 | 0/74 | 5/343 | 6/567 |
| Lesions per patient, median/mean (range) | 2/4.39 (1–18) | 4/4.83 (1–19) | 2/3.61 (1–18) | 4/6.09 (1–44) | 3/5.37 (1–44) |
| Disease volume, mean (range) (cc) | 1.581 (0.019–62.381) | 0.913 (0.008–21.640) | 1.259 (0.013–16.343) | 1.064 (0.002–48.750) | 1.151 (0.002–62.381) |
| Sphericity, mean (range) | 0.833 (0.531–0.914) | 0.827 (0.529–0.916) | 0.816 (0.504–0.899) | 0.805 (0.444–0.922) | 0.812 (0.444–0.922) |
| Histology top 3, N | Adenocarcinoma (84)<br>Other (2)<br>Unknown (1) | N/A<br>N/A<br>N/A | Other (65)<br>Adenocarcinoma (5)<br>Unknown (4) | Adenocarcinoma (337)<br>Other (8)<br>Bone (1) | Adenocarcinoma (426)<br>Other (75)<br>Unknown (5) |
| **Lung** |  |  |  |  |  |
| Patients, N | 222 | 253 | 248 | 600 | 1,323 |
| Lesions, N | 664 | 1,016 | 655 | 2,989 | 5,324 |
| Age, mean (range) | 65 (33–93) (N=201) | 66 (33–89) (N=151) | 65 (29–91) | 64 (32–91) | 65 (29–93) (N=1200) |
| Sex, N (M/F) | 106/116 | 114/139 | 120/128 | 274/326 | 614/709 |
| Lesions per patient, median/mean (range) | 2/2.99 (1–17) | 2/4.02 (1–21) | 2/2.64 (1–15) | 3/4.98 (1–48) | 2/4.02 (1–48) |
| Disease volume, mean (range) (cc) | 1.561 (0.032–25.541) | 0.893 (0.009–43.210) | 1.831 (0.007–40.484) | 1.108 (0.006–53.620) | 1.213 (0.006–53.620) |
| Sphericity, mean (range) | 0.849 (0.612–0.916) | 0.834 (0.545–0.922) | 0.831 (0.544–0.912) | 0.814 (0.561–0.948) | 0.827 (0.544–0.948) |
| Histology top 3 (N) | Adenocarcinoma (209)<br>Squamous Cell (8)<br>Other (3) | N/A<br>N/A<br>N/A | Adenocarcinoma (138)<br>Other (88)<br>Squamous Cell (20) | Adenocarcinoma (473)<br>Small Cell (31)<br>Carcinoma NOS (31) | Adenocarcinoma (980)<br>Other (91)<br>Squamous Cell (59) |
| **Melanoma** |  |  |  |  |  |
| Patients, N | 62 | 56 | 55 | 236 | 409 |
| Lesions, N | 249 | 287 | 148 | 1,312 | 1,996 |
| Age, mean (range) | 63 (20–91) (N=56) | 61 (27–85) (N=31) | 59 (23–81) | 61 (29–101) | 61 (20–101) (N=378) |
| Sex, N (M/F) | 40/22 | 40/16 | 37/18 | 147/89 | 264/145 |
| Lesions per patient, median/mean (range) | 2/4.02 (1–21) | 3/5.12 (1–27) | 2/2.69 (1–12) | 3/5.56 (1–51) | 3/4.88 (1–51) |
| Disease volume, mean (range) (cc) | 1.456 (0.030–38.727) | 1.362 (0.005–52.918) | 2.385 (0.005–14.842) | 1.326 (0.004–33.036) | 1.493 (0.004–52.918) |
| Sphericity, mean (range) | 0.847 (0.683–0.910) | 0.837 (0.654–0.906) | 0.824 (0.430–0.909) | 0.819 (0.502–0.922) | 0.825 (0.430–0.922) |
| Histology top 3, N | Melanoma (62) | Melanoma (56) | Melanoma (55) | Melanoma (236) | Melanoma (409) |

**TABLE S2.** Neurotransmitter receptors and transporters used to build the receptor cortical inter-regional similarity matrix. BP_ND_ = non-displaceable binding potential; V_T_ = tracer distribution volume; B_max_ = density (pmol/ml) converted from binding potential (5-HT) or distributional volume (GABA) using autoradiography-derived densities; SUVR = standard uptake value ratio. Values in parentheses (under #Participants) indicate number of females. Neurotransmitter receptor maps without citations refer to previously unpublished data. In those cases, contact information for the study principal investigator is provided in the Table. Asterisks indicate transporters.

| Receptor/<br>transporter | Neurotransmitter | Tracer | Measure | #Participants | Age (years) | References |
| --- | --- | --- | --- | --- | --- | --- |
| D<sub>1</sub> | dopamine | [<sup>11</sup>C]SCH23390 | BP<sub>ND</sub> | 13 (7) | 33 ± 13 | Kaller et al., 2017<sup>59</sup> |
| D<sub>2</sub> | dopamine | [<sup>11</sup>C]FLB-457 | BP<sub>ND</sub> | 37 (20) | 48.4 ± 16.9 | Smith et al., 2019<sup>101,112</sup> |
| D<sub>2</sub> | dopamine | [<sup>11</sup>C]FLB-457 | BP<sub>ND</sub> | 55 (29) | 32.5 ± 9.7 | Sandiego et al., 2015<sup>101,102,110,112,128</sup> |
| DAT* | dopamine | [<sup>123</sup>I]-FP-CIT | SUVR | 174 (65) | 61 ± 11 | Dukart et al., 2018<sup>28</sup> |
| NET* | norepinephrine | [<sup>11</sup>C]MRB | BP<sub>ND</sub> | 77 (27) | 33.4 ± 9.2 | Ding et al., 2010<sup>11,25,100</sup> |
| 5-HT<sub>1A</sub> | serotonin | [<sup>11</sup>C]WAY-100635 | BP<sub>ND</sub> | 35 (17) | 26.3 ± 5.2 | Savli et al., 2012<sup>104</sup> |
| 5-HT<sub>1B</sub> | serotonin | [<sup>11</sup>C]P943 | BP<sub>ND</sub> | 65 (16) | 33.7 ± 9.7 | Gallezot et al., 2010<sup>5,38,74,77,78,92,103</sup> |
| 5-HT<sub>1B</sub> | serotonin | [<sup>11</sup>C]P943 | BP<sub>ND</sub> | 23 (8) | 28.7 ± 7.0 | Savli et al., 2012<sup>104</sup> |
| 5-HT<sub>2A</sub> | serotonin | [<sup>11</sup>C]Cimbi-36 | B<sub>max</sub> | 29 (14) | 22.6 ± 2.7 | Beliveau et al., 2017<sup>12</sup> |
| 5-HT<sub>4</sub> | serotonin | [<sup>11</sup>C]SB207145 | B<sub>max</sub> | 59 (18) | 25.9 ± 5.3 | Beliveau et al., 2017<sup>12</sup> |
| 5-HT<sub>6</sub> | serotonin | [<sup>11</sup>C]GSK215083 | BP<sub>ND</sub> | 30 (0) | 36.6 ± 9.0 | Radhakrishnan et al., 2018<sup>95,96</sup> |
| 5-HTT* | serotonin | [<sup>11</sup>C]DASB | B<sub>max</sub> | 100 (71) | 25.1 ± 5.8 | Beliveau et al., 2017<sup>12</sup> |
| α<sub>4</sub>/β<sub>2</sub> | acetylcholine | [<sup>18</sup>F]flubatine | V<sub>T</sub> | 30 (10) | 33.5 ± 10.7 | Hillmer et al., 2016<sup>4,53</sup> |
| M<sub>1</sub> | acetylcholine | [<sup>11</sup>C]LSN3172176 | BP<sub>ND</sub> | 24 (11) | 40.5 ± 11.7 | Naganawa et al., 2021<sup>79</sup> |
| VACHT* | acetylcholine | [<sup>18</sup>F]FEOBV | SUVR | 4 (1) | 37 ± 10.2 | PI: L Tuominen & S Guimond |
| VACHT* | acetylcholine | [<sup>18</sup>F]FEOBV | SUVR | 18 (13) | 66.8 ± 6.8 | Aghourian et al., 2017<sup>2</sup> |
| VACHT* | acetylcholine | [<sup>18</sup>F]FEOBV | SUVR | 5 (1) | 68.3 ± 3.1 | Bedard et al., 2019<sup>10</sup> |
| VACHT* | acetylcholine | [<sup>18</sup>F]FEOBV | SUVR | 3 (3) | 66.6 ± 0.94 | PI: TW Schmitz & RN Spreng |
| mGluR<sub>5</sub> | glutamate | [<sup>11</sup>C]ABP688 | BP<sub>ND</sub> | 73 (48) | 19.9 ± 3.04 | Smart et al., 2019<sup>111</sup> |
| mGluR<sub>5</sub> | glutamate | [<sup>11</sup>C]ABP688 | BP<sub>ND</sub> | 22 (10) | 67.9 ± 9.6 | PI: P Rosa-Neto & E Kobayashi |
| mGluR<sub>5</sub> | glutamate | [<sup>11</sup>C]ABP688 | BP<sub>ND</sub> | 28 (13) | 33.1 ± 11.2 | DuBois et al., 2016<sup>27</sup> |
| GABA<sub>A/BZ</sub> | GABA | [<sup>11</sup>C]flumazenil | B<sub>max</sub> | 16 (9) | 26.6 ± 8 | Nørgaard et al., 2021<sup>84</sup> |
| H<sub>3</sub> | histamine | [<sup>11</sup>C]GSK189254 | V<sub>T</sub> | 8 (1) | 31.7 ± 9.0 | Gallezot et al., 2017<sup>39</sup> |
| CB<sub>1</sub> | cannabinoid | [<sup>11</sup>C]OMAR | V<sub>T</sub> | 77 (28) | 30.0 ± 8.9 | Normandin et al., 2015<sup>29,82,85,97</sup> |
| MOR | opioid | [<sup>11</sup>C]carfentanil | BP<sub>ND</sub> | 204 (72) | 32.3 ± 10.8 | Kantonen et al., 2020<sup>60</sup> |

## References

[1] Abraham, A., Pedregosa, F., Eickenberg, M., Gervais, P., Mueller, A., Kossaifi, J., Gramfort, A., Thirion, B., and Varoquaux, G. (2014). Machine learning for neuroimaging with scikit-learn. Frontiers in Neuroinformatics, 8:14.

[2] Aghourian, M., Legault-Denis, C., Soucy, J., Rosa-Neto, P., Gauthier, S., Kostikov, A., Gravel, P., and Bedard, M. (2017). Quantification of brain cholinergic denervation in alzheimer’s disease using pet imaging with [18 f]-feobv. Molecular psychiatry, 22(11):1531–1538.

[3] Amunts, K., Lepage, C., Borgeat, L., Mohlberg, H., Dickscheid, T., Rousseau, M.-É., Bludau, S., Bazin, P.-L., Lewis, L. B., Oros-Peusquens, A.-M., et al. (2013). Big-brain: an ultrahigh-resolution 3d human brain model. science, 340(6139):1472–1475.

[4] Baldassarri, S. R., Hillmer, A. T., Anderson, J. M., Jatlow, P., Nabulsi, N., Labaree, D., Cosgrove, K. P., O’Malley, S. S., Eissenberg, T., Krishnan-Sarin, S., et al. (2018). Use of electronic cigarettes leads to significant beta2-nicotinic acetylcholine receptor occupancy: evidence from a pet imaging study. Nicotine and Tobacco Research, 20(4):425–433.

[5] Baldassarri, S. R., Park, E., Finnema, S. J., Planeta, B., Nabulsi, N., Najafzadeh, S., Ropchan, J., Huang, Y., Hannestad, J., Maloney, K., et al. (2020). Inverse changes in raphe and cortical 5-ht1b receptor availability after acute tryptophan depletion in healthy human subjects. Synapse, 74(10):e22159.

[6] Bao, H., Ren, P., Liang, X., Lai, J., Bai, Y., Liu, Y., Lv, Z., Hu, J., Yan, Z., Wang, Z., et al. (2024). The spatial distribution of brain metastasis is determined by the heterogeneity of the brain microenvironment. Human Brain Mapping, 45(18):e70103.

[7] Bareche, Y., Venet, D., Ignatiadis, M., Aftimos, P., Piccart, M., Rothe, F., and Sotiriou, C. (2018). Unravelling triple-negative breast cancer molecular heterogeneity using an integrative multiomic analysis. Annals of Oncology, 29(4):895–902.

[8] Barrios, J., Porter, E., Capaldi, D. P., Upadhaya, T., Chen, W. C., Perks, J. R., Apte, A., Aristophanous, M., LoCastro, E., Hsu, D., et al. (2025). Multi-institutional atlas of brain metastases informs spatial modeling for precision imaging and personalized therapy. Nature Communications, 16(1):4536.

[9] Bazinet, V., Hansen, J. Y., and Misic, B. (2023). Towards a biologically annotated brain connectome. Nature Reviews Neuroscience, 24(12):747–760.

[10] Bedard, M.-A., Aghourian, M., Legault-Denis, C., Postuma, R. B., Soucy, J.-P., Gagnon, J.-F., Pelletier, A., and Montplaisir, J. (2019). Brain cholinergic alterations in idiopathic rem sleep behaviour disorder: a pet imaging study with 18f-feobv. Sleep medicine, 58:35–41.

[11] Belfort-DeAguiar, R., Gallezot, J.-D., Hwang, J. J., Elshafie, A., Yeckel, C. W., Chan, O., Carson, R. E., Ding, Y.-S., and Sherwin, R. S. (2018). Noradrenergic activity in the human brain: a mechanism supporting the defense against hypoglycemia. The Journal of Clinical Endocrinology & Metabolism, 103(6):2244–2252.

[12] Beliveau, V., Ganz, M., Feng, L., Ozenne, B., Højgaard, L., Fisher, P. M., Svarer, C., Greve, D. N., and Knudsen, G. M. (2017). A high-resolution in vivo atlas of the human brain’s serotonin system. Journal of Neuroscience, 37(1):120–128.

[13] Bergui, M., Castagno, D., D’Agata, F., Cicerale, A., Anselmino, M., Maria Ferrio, F., Giustetto, C., Halimi, F., Scaglione, M., and Gaita, F. (2015). Selective vulnerability of cortical border zone to microembolic infarct. Stroke, 46(7):1864–1869.

[14] Bookheimer, S. Y., Salat, D. H., Terpstra, M., Ances, B. M., Barch, D. M., Buckner, R. L., Burgess, G. C., Curtiss, S. W., Diaz-Santos, M., Elam, J. S., et al. (2019). The lifespan human connectome project in aging: an overview. Neuroimage, 185:335–348.

[15] Botstein, D., Cherry, J. M., Ashburner, M., Ball, C. A., Blake, J. A., Butler, H., Davis, A. P., Dolinski, K., Dwight, S. S., Eppig, J. T., et al. (2000). Gene ontology: tool for the unification of biology. Nat genet, 25(1):25–29.

[16] Brett, M., Markiewicz, C. J., Hanke, M., Côté, M.-A., Cipollini, B., McCarthy, P., Jarecka, D., Cheng, C. P., Halchenko, Y. O., Cottaar, M., Larson, E., Ghosh, S., Wassermann, D., Gerhard, S., Lee, G. R., Wang, H.-T., Kastman, E., Kaczmarzyk, J., Guidotti, R., Daniel, J., Duek, O., Rokem, A., Madison, C., Moloney, B., Morency, F. C., Goncalves, M., Markello, R., Riddell, C., Sólon, A., Burns, C., Millman, J., Gramfort, A., Leppäkangas, J., van den Bosch, J. J., Vincent, R. D., Braun, H., Subramaniam, K., Papadopoulos Orfanos, D., Van, A., Gorgolewski, K. J., Raamana, P. R., Klug, J., Nichols, B. N., Baker, E. M., Hayashi, S., Pinsard, B., Haselgrove, C., Hymers, M., Esteban, O., Koudoro, S., Pérez-García, F., Dockès, J., Oosterhof, N. N., Amirbekian, B., Nimmo-Smith, I., Nguyen, L., Reddigari, S., St-Jean, S., Panfilov, E., Garyfallidis, E., Varoquaux, G., Legarreta, J. H., Hahn, K. S., Waller, L., Hinds, O. P., Fauber, B., Roberts, J., Poline, J.-B., Stutters, J., Jordan, K., Cieslak, M., Moreno, M. E., Hrnčiar, T., Haenel, V., Schwartz, Y., Baratz, Z., Darwin, B. C., Thirion, B., Gauthier, C., Solovey, I., Gonzalez, I., Palasubramaniam, J., Lecher, J., Leinweber, K., Raktivan, K., Calábková, M., Fischer, P., Gervais, P., Gadde, S., Ballinger, T., Roos, T., Reddam, V. R., and freec84 (2022). nipy/nibabel:.

[17] Buie, J. J., Watson, L. S., Smith, C. J., and Sims-Robinson, C. (2019). Obesity-related cognitive impairment: The role of endothelial dysfunction. Neurobiology of Disease, 132:104580.

[18] Chaffer, C. L. and Weinberg, R. A. (2011). A perspective on cancer cell metastasis. Science, 331(6024):1559– 1564.

[19] Chen, G., Chakravarti, N., Aardalen, K., Lazar, A. J., Tetzlaff, M. T., Wubbenhorst, B., Kim, S.-B., Kopetz, S., Ledoux, A. A., Gopal, Y. V., et al. (2014). Molecular profiling of patient-matched brain and extracranial melanoma metastases implicates the pi3k pathway as a therapeutic target. Clinical Cancer Research, 20(21):5537–5546.

[20] Clark, K., Vendt, B., Smith, K., Freymann, J., Kirby, J., Koppel, P., Moore, S., Phillips, S., Maffitt, D., Pringle, M., et al. (2013). The cancer imaging archive (tcia): maintaining and operating a public information repository. Journal of Digital Imaging, 26(6):1045–1057.

[21] Davies, M. A. (2012). The role of the pi3k-akt pathway in melanoma. The Cancer Journal, 18(2):142–147.

[22] Delattre, J. Y., Krol, G., Thaler, H. T., and Posner, J. B. (1988). Distribution of brain metastases. Archives of Neurology, 45(7):741–744.

[23] Dianat-Moghadam, H., Azizi, M., Eslami-S, Z., Cortés-Hernández, L. E., Heidarifard, M., Nouri, M., and Alix-Panabières, C. (2020). The role of circulating tumor cells in the metastatic cascade: biology, technical challenges, and clinical relevance. Cancers, 12(4):867.

[24] Diedrichsen, J., Balsters, J. H., Flavell, J., Cussans, E., and Ramnani, N. (2009). A probabilistic mr atlas of the human cerebellum. Neuroimage, 46(1):39–46.

[25] Ding, Y.-S., Singhal, T., Planeta-Wilson, B., Gallezot, J.-D., Nabulsi, N., Labaree, D., Ropchan, J., Henry, S., Williams, W., Carson, R. E., et al. (2010). Pet imaging of the effects of age and cocaine on the norepinephrine transporter in the human brain using (s, s)-[11c] omethylreboxetine and hrrt. Synapse, 64(1):30–38.

[26] Dray, S. (2011). A new perspective about moran’s coefficient: spatial autocorrelation as a linear regression problem. moran. Geographical Analysis, 43(2):127–141.

[27] DuBois, J. M., Rousset, O. G., Rowley, J., Porras-Betancourt, M., Reader, A. J., Labbe, A., Massarweh, G., Soucy, J.-P., Rosa-Neto, P., and Kobayashi, E. (2016). Characterization of age/sex and the regional distribution of mglur5 availability in the healthy human brain measured by high-resolution [11 c] abp688 pet. European journal of nuclear medicine and molecular imaging, 43(1):152–162.

[28] Dukart, J., Holiga, Š., Chatham, C., Hawkins, P., Forsyth, A., McMillan, R., Myers, J., Lingford-Hughes, A. R., Nutt, D. J., Merlo-Pich, E., et al. (2018). Cerebral blood flow predicts differential neurotransmitter activity. Scientific reports, 8(1):1–11.

[29] D’Souza, D. C., Cortes-Briones, J. A., Ranganathan, M., Thurnauer, H., Creatura, G., Surti, T., Planeta, B., Neumeister, A., Pittman, B., Normandin, M. D., et al. (2016). Rapid changes in cannabinoid 1 receptor availability in cannabis-dependent male subjects after abstinence from cannabis. Biological psychiatry: cognitive neuroscience and neuroimaging, 1(1):60–67.

[30] Ewing, J. (1928). Neoplastic diseases: a treatise on tumors. Wb saunders.

[31] Farahani, A., Hansen, J. Y., Bazinet, V., Shafiei, G., Collins, D. L., Dadar, M., Kalra, S., Dagher, A., and Misic, B. (2025a). Network spreading and local biological vulnerability in amyotrophic lateral sclerosis. Communications Biology, 8(1):1153.

[32] Farahani, A., Liu, Z.-Q., Ceballos, E. G., Hansen, J. Y., Wennberg, K., Zeighami, Y., Dadar, M., Gauthier, C. J., Dagher, A., and Misic, B. (2025b). Mapping cerebral blood perfusion and its links to multi-scale brain organization across the human lifespan. Plos Biology, 23(7):e3003277.

[33] Farahani, A., Liu, Z.-Q., Morys, F., Moqadam, R., Zeighami, Y., Dadar, M., Dagher, A., and Misic, B. (2026). Aging and metabolism contribute separately to brain–body health. Plos Biology, 24(6):e3003856.

[34] Fidler, I. J. (1978). Tumor heterogeneity and the biology of cancer invasion and metastasis. Cancer Research, 38(9):2651–2660.

[35] Follain, G., Herrmann, D., Harlepp, S., Hyenne, V., Osmani, N., Warren, S. C., Timpson, P., and Goetz, J. G. (2020). Fluids and their mechanics in tumour transit: shaping metastasis. Nature Reviews Cancer, 20(2):107– 124.

[36] Follain, G., Osmani, N., Azevedo, A. S., Allio, G., Mercier, L., Karreman, M. A., Solecki, G., Leòn, M. J. G., Lefebvre, O., Fekonja, N., et al. (2018). Hemodynamic forces tune the arrest, adhesion, and extravasation of circulating tumor cells. Developmental Cell, 45(1):33–52.

[37] Fulcher, B. D., Arnatkeviciute, A., and Fornito, A. (2021). Overcoming false-positive gene-category enrichment in the analysis of spatially resolved transcriptomic brain atlas data. Nature Communications, 12(1):2669.

[38] Gallezot, J.-D., Nabulsi, N., Neumeister, A., Planeta-Wilson, B., Williams, W. A., Singhal, T., Kim, S., Maguire, R. P., McCarthy, T., Frost, J. J., et al. (2010). Kinetic modeling of the serotonin 5-ht1b receptor radioligand [11c] p943 in humans. Journal of Cerebral Blood Flow & Metabolism, 30(1):196–210.

[39] Gallezot, J.-D., Planeta, B., Nabulsi, N., Palumbo, D., Li, X., Liu, J., Rowinski, C., Chidsey, K., Labaree, D., Ropchan, J., et al. (2017). Determination of receptor occupancy in the presence of mass dose:[11c] gsk189254 pet imaging of histamine h3 receptor occupancy by pf-03654746. Journal of Cerebral Blood Flow & Metabolism, 37(3):1095–1107.

[40] Gavrilovic, I. T. and Posner, J. B. (2005). Brain metastases: epidemiology and pathophysiology. Journal of Neuro-oncology, 75(1):5–14.

[41] Gerstberger, S., Jiang, Q., and Ganesh, K. (2023). Metastasis. Cell, 186(8):1564–1579.

[42] Glasser, M. F., Smith, S. M., Marcus, D. S., Andersson, J. L., Auerbach, E. J., Behrens, T. E., Coalson, T. S., Harms, M. P., Jenkinson, M., Moeller, S., et al. (2016). The human connectome project’s neuroimaging approach. Nature Neuroscience, 19(9):1175–1187.

[43] Glasser, M. F., Sotiropoulos, S. N., Wilson, J. A., Coalson, T. S., Fischl, B., Andersson, J. L., Xu, J., Jbabdi, S., Webster, M., Polimeni, J. R., et al. (2013). The minimal preprocessing pipelines for the human connectome project. Neuroimage, 80:105–124.

[44] Goncalves, P. H., Peterson, S. L., Vigneau, F. D., Shore, R. D., Quarshie, W. O., Islam, K., Schwartz, A. G., Wozniak, A. J., and Gadgeel, S. M. (2016). Risk of brain metastases in patients with nonmetastatic lung cancer: Analysis of the metropolitan detroit surveillance, epidemiology, and end results (seer) data. Cancer, 122(12):1921–1927.

[45] Guerrini, V. and Gennaro, M. L. (2019). Foam cells: one size doesn’t fit all. Trends in Immunology, 40(12):1163–1179.

[46] Gupta, G. P. and Massagué, J. (2006). Cancer metastasis: building a framework. Cell, 127(4):679–695.

[47] Gustafson, D., Karlsson, C., Skoog, I., Rosengren, L., Lissner, L., and Blennow, K. (2007). Mid-life adiposity factors relate to blood–brain barrier integrity in late life. Journal of Internal Medicine, 262(6):643–650.

[48] Hansen, J. Y., Shafiei, G., Markello, R. D., Smart, K., Cox, S. M., Nørgaard, M., Beliveau, V., Wu, Y., Gallezot, J.-D., Aumont, É., et al. (2022). Mapping neurotransmitter systems to the structural and functional organization of the human neocortex. Nature neuroscience, 25(11):1569–1581.

[49] Hansen, J. Y., Shafiei, G., Voigt, K., Liang, E. X., Cox, S. M., Leyton, M., Jamadar, S. D., and Misic, B. (2023). Integrating multimodal and multiscale connectivity blueprints of the human cerebral cortex in health and disease. Plos Biology, 21(9):e3002314.

[50] Harms, M. P., Somerville, L. H., Ances, B. M., Andersson, J., Barch, D. M., Bastiani, M., Bookheimer, S. Y., Brown, T. B., Buckner, R. L., Burgess, G. C., et al. (2018). Extending the human connectome project across ages: Imaging protocols for the lifespan development and aging projects. Neuroimage, 183:972–984.

[51] Harris, C. R., Millman, K. J., Van Der Walt, S. J., Gommers, R., Virtanen, P., Cournapeau, D., Wieser, E., Taylor, J., Berg, S., Smith, N. J., et al. (2020). Array programming with numpy. Nature, 585(7825):357–362.

[52] Hawrylycz, M. J., Lein, E. S., Guillozet-Bongaarts, A. L., Shen, E. H., Ng, L., Miller, J. A., Van De Lagemaat, L. N., Smith, K. A., Ebbert, A., Riley, Z. L., et al. (2012). An anatomically comprehensive atlas of the adult human brain transcriptome. Nature, 489(7416):391–399.

[53] Hillmer, A. T., Esterlis, I., Gallezot, J.-D., Bois, F., Zheng, M.-Q., Nabulsi, N., Lin, S.-F., Papke, R., Huang, Y., Sabri, O., et al. (2016). Imaging of cerebral α4β2* nicotinic acetylcholine receptors with (-)-[18f] flubatine pet: Implementation of bolus plus constant infusion and sensitivity to acetylcholine in human brain. Neuroimage, 141:71–80.

[54] Hunter, J. D. (2007). Matplotlib: A 2d graphics environment. Computing in Science & Engineering, 9(03):90–95.

[55] Hwang, T.-L., Close, T. P., Grego, J. M., Brannon, W. L., and Gonzales, F. (1996). Predilection of brain metastasis in gray and white matter junction and vascular border zones. Cancer: Interdisciplinary International Journal of the American Cancer Society, 77(8):1551–1555.

[56] Jamadar, S. D., Ward, P. G., Close, T. G., Fornito, A., Premaratne, M., O’Brien, K., Stäb, D., Chen, Z., Shah, N. J., and Egan, G. F. (2020). Simultaneous bold-fmri and constant infusion fdg-pet data of the resting human brain. Scientific data, 7(1):363.

[57] Ji, J. L., Demšar, J., Fonteneau, C., Tamayo, Z., Pan, L., Kraljič, A., Matkovič, A., Purg, N., Helmer, M., Warrington, S., et al. (2023). Qunex—an integrative platform for reproducible neuroimaging analytics. Frontiers in Neuroinformatics, 17:1104508.

[58] Juttukonda, M. R., Li, B., Almaktoum, R., Stephens, K. A., Yochim, K. M., Yacoub, E., Buckner, R. L., and Salat, D. H. (2021). Characterizing cerebral hemodynamics across the adult lifespan with arterial spin labeling mri data from the human connectome project-aging. Neuroimage, 230:117807.

[59] Kaller, S., Rullmann, M., Patt, M., Becker, G.-A., Luthardt, J., Girbardt, J., Meyer, P. M., Werner, P., Barthel, H., Bresch, A., et al. (2017). Test–retest measurements of dopamine d 1-type receptors using simultaneous pet/mri imaging. European journal of nuclear medicine and molecular imaging, 44(6):1025–1032.

[60] Kantonen, T., Karjalainen, T., Isojärvi, J., Nuutila, P., Tuisku, J., Rinne, J., Hietala, J., Kaasinen, V., Kalliokoski, K., Scheinin, H., et al. (2020). Interindividual variability and lateralization of µ-opioid receptors in the human brain. NeuroImage, 217:116922.

[61] Kircher, D. A., Silvis, M. R., Cho, J. H., and Holmen, S. L. (2016). Melanoma brain metastasis: mechanisms, models, and medicine. International Journal of Molecular Sciences, 17(9):1468.

[62] Kirk, T. F., McConnell, F. A. K., Toner, J., Craig, M. S., Carone, D., Li, X., Suzuki, Y., Coalson, T. S., Harms, M. P., Glasser, M. F., and Chappell, M. A. (2025). Arterial spin labelling perfusion mri analysis for the human connectome project lifespan ageing and development studies. Imaging Neuroscience, 3.

[63] Kirst, C., Skriabine, S., Vieites-Prado, A., Topilko, T., Bertin, P., Gerschenfeld, G., Verny, F., Topilko, P., Michalski, N., Tessier-Lavigne, M., et al. (2020). Mapping the fine-scale organization and plasticity of the brain vasculature. Cell, 180(4):780–795.

[64] Kyeong, S., Cha, Y. J., Ahn, S. G., Suh, S. H., Son, E. J., and Ahn, S. J. (2017). Subtypes of breast cancer show different spatial distributions of brain metastases. PLoS One, 12(11):e0188542.

[65] Lambert, A. W., Pattabiraman, D. R., and Weinberg, R. A. (2017). Emerging biological principles of metastasis. Cell, 168(4):670–691.

[66] Lasocki, A., Khoo, C., Lau, P. K., Kok, D. L., and McArthur, G. A. (2020). High-resolution mri demonstrates that more than 90% of small intracranial melanoma metastases develop in close relationship to the leptomeninges. Neuro-oncology, 22(3):423–432.

[67] Liotta, L. and Petricoin, E. (2000). Molecular profiling of human cancer. Nature Reviews Genetics, 1(1):48–56.

[68] Liu, Z.-Q., Bazinet, V., Hansen, J. Y., Milisav, F., Luppi, A. I., Ceballos, E. G., Farahani, A., Suarez, L. E., Shafiei, G., Markello, R. D., et al. (2025). netneurotools: a trainee-oriented approach to network neuroscience. bioRxiv, pages 2025–09.

[69] Lyu, Q., Yuan, H., Lin, Z., Barcus, R., Hudson, J., Jiang, Y., Kim, J., and Whitlow, C. T. (2025). A large scale multi-dataset investigation of brain metastases distribution based on primary cancer type. BMC Cancer, 25(1):1821.

[70] Marcus, D. S., Harms, M. P., Snyder, A. Z., Jenkinson, M., Wilson, J. A., Glasser, M. F., Barch, D. M., Archie, K. A., Burgess, G. C., Ramaratnam, M., et al. (2013). Human connectome project informatics: quality control, database services, and data visualization. Neuroimage, 80:202–219.

[71] Markello, R. D., Arnatkeviciute, A., Poline, J.-B., Fulcher, B. D., Fornito, A., and Misic, B. (2021). Standardizing workflows in imaging transcriptomics with the abagen toolbox. eLife, 10:e72129.

[72] Markello, R. D., Hansen, J. Y., Liu, Z.-Q., Bazinet, V., Shafiei, G., Suárez, L. E., Blostein, N., Seidlitz, J., Baillet, S., Satterthwaite, T. D., et al. (2022). Neuromaps: structural and functional interpretation of brain maps. Nature Methods, 19(11):1472–1479.

[73] Massagué, J. and Obenauf, A. C. (2016). Metastatic colonization by circulating tumour cells. Nature, 529(7586):298–306.

[74] Matuskey, D., Bhagwagar, Z., Planeta, B., Pittman, B., Gallezot, J.-D., Chen, J., Wanyiri, J., Najafzadeh, S., Ropchan, J., Geha, P., et al. (2014). Reductions in brain 5-ht1b receptor availability in primarily cocaine-dependent humans. Biological psychiatry, 76(10):816–822.

[75] McKinney, W. et al. (2010). Data structures for statistical computing in python. SciPy, 445(1):51–56.

[76] Mirzaeva, M., Mirzayev, A., Iriskulov, B., Azimova, S. B., Nizamkhodjaev, S., and Asrorov, A. M. (2026). Endothelial dysfunction determines vascular mechanisms of metastatic progression in breast cancer. Discover Oncology.

[77] Murrough, J. W., Czermak, C., Henry, S., Nabulsi, N., Gallezot, J.-D., Gueorguieva, R., Planeta-Wilson, B., Krystal, J. H., Neumaier, J. F., Huang, Y., et al. (2011a). The effect of early trauma exposure on serotonin type 1b receptor expression revealed by reduced selective radioligand binding. Archives of general psychiatry, 68(9):892–900.

[78] Murrough, J. W., Henry, S., Hu, J., Gallezot, J.-D., Planeta-Wilson, B., Neumaier, J. F., and Neumeister, A. (2011b). Reduced ventral striatal/ventral pallidal serotonin 1b receptor binding potential in major depressive disorder. Psychopharmacology, 213:547–553.

[79] Naganawa, M., Nabulsi, N., Henry, S., Matuskey, D., Lin, S.-F., Slieker, L., Schwarz, A. J., Kant, N., Jesudason, C., Ruley, K., et al. (2021). First-in-human assessment of 11c-lsn3172176, an m1 muscarinic acetylcholine receptor pet radiotracer. Journal of Nuclear Medicine, 62(4):553–560.

[80] Nayak, L., Lee, E. Q., and Wen, P. Y. (2012). Epidemiology of brain metastases. Current Oncology Reports, 14(1):48–54.

[81] Network, C. G. A. R. et al. (2014). Comprehensive molecular profiling of lung adenocarcinoma. Nature, 511(7511):543.

[82] Neumeister, A., Normandin, M. D., Murrough, J. W., Henry, S., Bailey, C. R., Luckenbaugh, D. A., Tuit, K., Zheng, M.-Q., Galatzer-Levy, I. R., Sinha, R., et al. (2012). Positron emission tomography shows elevated cannabinoid cb 1 receptor binding in men with alcohol dependence. Alcoholism: Clinical and Experimental Research, 36(12):2104–2109.

[83] Niessner, H., Schmitz, J., Tabatabai, G., Schmid, A. M., Calaminus, C., Sinnberg, T., Weide, B., Eigentler, T. K., Garbe, C., Schittek, B., et al. (2016). Pi3k pathway inhibition achieves potent antitumor activity in melanoma brain metastases in vitro and in vivo. Clinical Cancer Research, 22(23):5818–5828.

[84] Nørgaard, M., Beliveau, V., Ganz, M., Svarer, C., Pinborg, L. H., Keller, S. H., Jensen, P. S., Greve, D. N., and Knudsen, G. M. (2021). A high-resolution in vivo atlas of the human brain’s benzodiazepine binding site of gabaa receptors. NeuroImage, 232:117878.

[85] Normandin, M. D., Zheng, M.-Q., Lin, K.-S., Mason, N. S., Lin, S.-F., Ropchan, J., Labaree, D., Henry, S., Williams, W. A., Carson, R. E., et al. (2015). Imaging the cannabinoid cb1 receptor in humans with [11c] omar: assessment of kinetic analysis methods, test–retest reproducibility, and gender differences. Journal of Cerebral Blood Flow & Metabolism, 35(8):1313–1322.

[86] Osman, M. A. and Hennessy, B. T. (2015). Obesity correlation with metastases development and response to first-line metastatic chemotherapy in breast cancer. Clinical Medicine Insights: Oncology, 9:CMO–S32812.

[87] Paget, S. (1989). The distribution of secondary growths in cancer of the breast. Cancer Metastasis Rev, 8:98–101.

[88] Paquola, C., Royer, J., Lewis, L. B., Lepage, C., Glatard, T., Wagstyl, K., DeKraker, J., Toussaint, P.-J., Valk, S. L., Collins, L., et al. (2021). The bigbrainwarp toolbox for integration of bigbrain 3d histology with multimodal neuroimaging. Elife, 10:e70119.

[89] Paquola, C., Vos De Wael, R., Wagstyl, K., Bethlehem, R. A., Hong, S.-J., Seidlitz, J., Bullmore, E. T., Evans, A. C., Misic, B., Margulies, D. S., et al. (2019). Microstructural and functional gradients are increasingly dissociated in transmodal cortices. Plos Biology, 17(5):e3000284.

[90] Park, W., Lee, J.-S., Gao, G., Kim, B. S., and Cho, D.-W. (2023). 3d bioprinted multilayered cerebrovascular conduits to study cancer extravasation mechanism related with vascular geometry. Nature Communications, 14(1):7696.

[91] Pathak, S., Palkhi, E., Dave, R., White, A., Pandanaboyana, S., Prasad, K. R., Lodge, J. P. A., and Toogood, G. J. (2016). Relationship between primary colorectal tumour and location of colorectal liver metastases. ANZ Journal of Surgery, 86(5):408–410.

[92] Pittenger, C., Adams Jr, T. G., Gallezot, J.-D., Crowley, M. J., Nabulsi, N., Ropchan, J., Gao, H., Kichuk, S. A., Simpson, R., Billingslea, E., et al. (2016). Ocd is associated with an altered association between sensorimotor gating and cortical and subcortical 5-ht1b receptor binding. Journal of affective disorders, 196:87–96.

[93] Prasad, S., Sajja, R. K., Naik, P., and Cucullo, L. (2014). Diabetes mellitus and blood-brain barrier dysfunction: an overview. Journal of Pharmacovigilance, 2(2):125.

[94] Qiu, Y., Chen, A., Yu, R., Llevenes, P., Seen, M., Ko, N. Y., Monti, S., and Denis, G. V. (2025). Insulin resistance increases tnbc aggressiveness and brain metastasis via adipocyte-derived exosomes. Molecular Cancer Research, 23(6):567–578.

[95] Radhakrishnan, R., Matuskey, D., Nabulsi, N., Gaiser, E., Gallezot, J.-D., Henry, S., Planeta, B., Lin, S.-f., Ropchan, J., Huang, Y., et al. (2020). In vivo 5-ht6 and 5-ht2a receptor availability in antipsychotic treated schizophrenia patients vs. unmedicated healthy humans measured with [11c] gsk215083 pet. Psychiatry Research: Neuroimaging, 295:111007.

[96] Radhakrishnan, R., Nabulsi, N., Gaiser, E., Gallezot, J.-D., Henry, S., Planeta, B., Lin, S.-f., Ropchan, J., Williams, W., Morris, E., et al. (2018). Age-related change in 5-ht6 receptor availability in healthy male volunteers measured with 11c-gsk215083 pet. Journal of Nuclear Medicine, 59(9):1445–1450.

[97] Ranganathan, M., Cortes-Briones, J., Radhakrishnan, R., Thurnauer, H., Planeta, B., Skosnik, P., Gao, H., Labaree, D., Neumeister, A., Pittman, B., et al. (2016). Reduced brain cannabinoid receptor availability in schizophrenia. Biological psychiatry, 79(12):997–1005.

[98] Redmer, T. (2018). Deciphering mechanisms of brain metastasis in melanoma-the gist of the matter. Molecular Cancer, 17(1):106.

[99] Rubinov, M. and Sporns, O. (2010). Complex network measures of brain connectivity: uses and interpretations. Neuroimage, 52(3):1059–1069.

[100] Sanchez-Rangel, E., Gallezot, J.-D., Yeckel, C. W., Lam, W., Belfort-DeAguiar, R., Chen, M.-K., Carson, R. E., Sherwin, R., and Hwang, J. J. (2020). Norepinephrine transporter availability in brown fat is reduced in obesity: a human pet study with [11c] mrb. International Journal of Obesity, 44(4):964–967.

[101] Sandiego, C. M., Gallezot, J.-D., Lim, K., Ropchan, J., Lin, S.-f., Gao, H., Morris, E. D., and Cosgrove, K. P. (2015). Reference region modeling approaches for amphetamine challenge studies with [11c] flb 457 and pet. Journal of Cerebral Blood Flow & Metabolism, 35(4):623– 629.

[102] Sandiego, C. M., Matuskey, D., Lavery, M., McGovern, E., Huang, Y., Nabulsi, N., Ropchan, J., Picciotto, M. R., Morris, E. D., McKee, S. A., et al. (2018). The effect of treatment with guanfacine, an alpha2 adrenergic agonist, on dopaminergic tone in tobacco smokers: an [11c] flb457 pet study. Neuropsychopharmacology, 43(5):1052–1058.

[103] Saricicek, A., Chen, J., Planeta, B., Ruf, B., Subramanyam, K., Maloney, K., Matuskey, D., Labaree, D., Deserno, L., Neumeister, A., et al. (2015). Test–retest reliability of the novel 5-ht 1b receptor pet radioligand [11 c] p943. European journal of nuclear medicine and molecular imaging, 42:468–477.

[104] Savli, M., Bauer, A., Mitterhauser, M., Ding, Y.-S., Hahn, A., Kroll, T., Neumeister, A., Haeusler, D., Ungersboeck, J., Henry, S., et al. (2012). Normative database of the serotonergic system in healthy subjects using multitracer pet. Neuroimage, 63(1):447–459.

[105] Schaefer, A., Kong, R., Gordon, E. M., Laumann, T. O., Zuo, X.-N., Holmes, A. J., Eickhoff, S. B., and Yeo, B. T. (2018). Local-global parcellation of the human cerebral cortex from intrinsic functional connectivity mri. Cerebral Cortex, 28(9):3095–3114.

[106] Schouten, L. J., Rutten, J., Huveneers, H. A., and Twijnstra, A. (2002). Incidence of brain metastases in a cohort of patients with carcinoma of the breast, colon, kidney, and lung and melanoma. Cancer, 94(10):2698–2705.

[107] Schroeder, T., Bittrich, P., Kuhne, J., Noebel, C., Leischner, H., Fiehler, J., Schroeder, J., Schoen, G., and Gellißen, S. (2020). Mapping distribution of brain metastases: does the primary tumor matter? Journal of Neurooncology, 147(1):229–235.

[108] Seabold, S. and Perktold, J. (2010). statsmodels: Econometric and statistical modeling with python. In 9th Python in Science Conference.

[109] Shafiei, G., Markello, R. D., Makowski, C., Talpalaru, A., Kirschner, M., Devenyi, G. A., Guma, E., Hagmann, P., Cashman, N. R., Lepage, M., et al. (2020). Spatial patterning of tissue volume loss in schizophrenia reflects brain network architecture. Biological Psychiatry, 87(8):727–735.

[110] Slifstein, M., Van De Giessen, E., Van Snellenberg, J., Thompson, J. L., Narendran, R., Gil, R., Hackett, E., Girgis, R., Ojeil, N., Moore, H., et al. (2015). Deficits in prefrontal cortical and extrastriatal dopamine release in schizophrenia: a positron emission tomographic functional magnetic resonance imaging study. JAMA psychiatry, 72(4):316–324.

[111] Smart, K., Cox, S. M., Scala, S. G., Tippler, M., Jaworska, N., Boivin, M., Séguin, J. R., Benkelfat, C., and Leyton, M. (2019). Sex differences in [11 c] abp688 binding: a positron emission tomography study of mglu5 receptors. European journal of nuclear medicine and molecular imaging, 46(5):1179–1183.

[112] Smith, C. T., Crawford, J. L., Dang, L. C., Seaman, K. L., San Juan, M. D., Vijay, A., Katz, D. T., Matuskey, D., Cowan, R. L., Morris, E. D., et al. (2019). Partial-volume correction increases estimated dopamine d2-like receptor binding potential and reduces adult age differences. Journal of Cerebral Blood Flow & Metabolism, 39(5):822– 833.

[113] Somerville, L. H., Bookheimer, S. Y., Buckner, R. L., Burgess, G. C., Curtiss, S. W., Dapretto, M., Elam, J. S., Gaffrey, M. S., Harms, M. P., Hodge, C., et al. (2018). The lifespan human connectome project in development: A large-scale study of brain connectivity development in 5–21 year olds. Neuroimage, 183:456–468.

[114] Taheri, T., Farahani, A., Liu, Z.-Q., Ceballos, E. G., Harroud, A., Dagher, A., and Misic, B. (2026). Spatial organization of aqp4 channels in the human brain: links with perfusion, edema, and disease vulnerability. bioRxiv, pages 2026–02.

[115] Takano, K., Kinoshita, M., Takagaki, M., Sakai, M., Tateishi, S., Achiha, T., Hirayama, R., Nishino, K., Uchida, J., Kumagai, T., et al. (2016). Different spatial distributions of brain metastases from lung cancer by histological subtype and mutation status of epidermal growth factor receptor. Neuro-oncology, 18(5):716–724.

[116] Tehranian, C., Fankhauser, L., Harter, P. N., Ratcliffe, C. D., Zeiner, P. S., Messmer, J. M., Hoffmann, D. C., Frey, K., Westphal, D., Ronellenfitsch, M. W., et al. (2022). The pi3k/akt/mtor pathway as a preventive target in melanoma brain metastasis. Neuro-oncology, 24(2):213–225.

[117] Van Der Walt, S., Colbert, S. C., and Varoquaux, G. (2011). The numpy array: a structure for efficient numerical computation. Computing in Science & Engineering, 13(2):22–30.

[118] Van Dyken, P. and Lacoste, B. (2018). Impact of metabolic syndrome on neuroinflammation and the blood–brain barrier. Frontiers in Neuroscience, 12:930.

[119] Van Essen, D. C., Smith, S. M., Barch, D. M., Behrens, T. E., Yacoub, E., Ugurbil, K., Consortium, W.-M. H., et al. (2013). The wu-minn human connectome project: an overview. Neuroimage, 80:62–79.

[120] Virtanen, P., Gommers, R., Oliphant, T. E., Haberland, M., Reddy, T., Cournapeau, D., Burovski, E., Peterson, P., Weckesser, W., Bright, J., et al. (2020). Scipy 1.0: fundamental algorithms for scientific computing in python. Nature Methods, 17(3):261–272.

[121] Vo, A., Tremblay, C., Rahayel, S., Shafiei, G., Hansen, J. Y., Yau, Y., Misic, B., and Dagher, A. (2023). Network connectivity and local transcriptomic vulnerability underpin cortical atrophy progression in parkinson’s disease. Neuroimage: Clinical, 40:103523.

[122] Voigt, K., Liang, E. X., Misic, B., Ward, P. G., Egan, G. F., and Jamadar, S. D. (2023). Metabolic and functional connectivity provide unique and complementary insights into cognition-connectome relationships. Cerebral Cortex, 33(4):1476–1488.

[123] Wagner, H. H. and Dray, S. (2015). Generating spatially constrained null models for irregularly spaced data using m oran spectral randomization methods. Methods in Ecology and Evolution, 6(10):1169–1178.

[124] Waskom, M. L. (2021). seaborn: statistical data visualization. Journal of Open Source Software, 6(60):3021.

[125] Wigmore, S. J., Madhavan, K., Redhead, D. N., Currie, E. J., and Garden, O. J. (2000). Distribution of colorectal liver metastases in patients referred for hepatic resection. Cancer: Interdisciplinary International Journal of the American Cancer Society, 89(2):285–287.

[126] Yong, S. W., Bang, O. Y., Lee, P. H., and Li, W. Y. (2006). Internal and cortical border-zone infarction: clinical and diffusion-weighted imaging features. Stroke, 37(3):841– 846.

[127] Yu, Y. P. and Tan, L. (2016). The vulnerability of vessels involved in the role of embolism and hypoperfusion in the mechanisms of ischemic cerebrovascular diseases. BioMed Research International, 2016(1):8531958.

[128] Zakiniaeiz, Y., Hillmer, A. T., Matuskey, D., Nabulsi, N., Ropchan, J., Mazure, C. M., Picciotto, M. R., Huang, Y., McKee, S. A., Morris, E. D., et al. (2019). Sex differences in amphetamine-induced dopamine release in the dorsolateral prefrontal cortex of tobacco smokers. Neuropsychopharmacology, 44(13):2205–2211.

